# An endothelial-specific TIE2-L914F mouse model reveals cellular mechanisms of venous malformation progression and recapitulates the patients’ response to targeted therapy

**DOI:** 10.64898/2026.10.29.762550

**Authors:** Lindsay J Bischoff, Sandra Schrenk, Andrew Barney, Kara Soroko, Chhiring Sherpa, Michael Simons, Elisa Boscolo

## Abstract

TIE2 p.L914F is the most common driver mutation of sporadic venous malformation (VM), a disease characterized by massively enlarged veins. The study of the molecular and cellular mechanisms contributing to TIE2-driven VM has been hindered by the lack of a genetic *in vivo* model. We generated a mouse system in which TIE2^L914F^ is expressed postnatally in endothelial cells (EC). Mutant mice develop visible lesions with dilated veins with irregular smooth muscle cell coverage, histologically mimicking patients’ VM.

With the use of an mCherry reporter we demonstrate that lesions form within the venous vasculature, and mutant ECs undergo clonal expansion fueled by a biphasic mechanism consisting of an early burst of proliferation followed by cell size increase. Targeted treatment with rapamycin partially reduces lesion growth by targeting increased proliferation and prolongs mouse survival, demonstrating this genetic murine model recapitulates the patients’ therapeutic response. Together, these findings provide new cellular and mechanistic insights into TIE2 p.L914F-driven venous malformation and establish a genetic platform for preclinical therapeutic studies.

## Introduction

Vascular anomalies encompass a range of disorders that involve abnormally structured and dysfunctional blood or lymphatic vessels. Under the classification proposed by the International Society for the Study of Vascular Anomalies (ISSVA), they are divided into two major categories: vascular tumors and vascular malformations (1). Vascular malformations are thought to result from errors in normal vascular development and often arise early in life, congenitally or during childhood. The most common type of vascular malformation is venous malformation (VM), which has an estimated incidence of 1-2 per 10,000 births (2, 3). VMs consist of abnormally dilated slow-flow veins that are often found in the skin, subcutaneous tissues, or muscle of the extremities or head and neck region (4). VMs cause significant morbidity for patients, including chronic pain, severe disfigurement, functional impairment of the affected tissue, and possibly life-threatening complications such as bleeding and coagulopathy. Together, these reduce the quality of life of VM patients (5–7).

VM is rarely inherited and it most often manifests as sporadic disease consisting of unifocal or multifocal lesions (8, 9). The majority of sporadic VM cases are driven by somatic activating mutations in the *TEK* gene, which encodes the TIE2 receptor tyrosine kinase (8–11). The most common TIE2 mutation (occurring in up to 80% of TIE2-mutant VM cases) is the p.L914F missense mutation, which causes ligand-independent hyperactivation of the receptor (9, 12–14), and is specifically found within lesional endothelial cells (EC) (15). The TIE2 p.L914F mutation has never been identified in hereditary, germline cases of VM (10) and was shown to be embryonically lethal when expressed in the germline in a transgenic *TIE2^L914F^* mouse model (16). Therefore, TIE2 p.L914F mutation is thought to cause VM when it occurs in the mosaic or somatic condition, which is similar for other developmental disorders (17, 18). We have recently shown that constitutive but mosaic expression of mutant TIE2 during mouse development is only partially embryonically lethal and surviving mice develop a VM-like phenotype (19).

The molecular and cellular mechanisms that drive formation of TIE2 p.L914F-driven VM have been revealed by *in vitro* studies (9, 12, 14, 20, 21) and mouse xenografts using human TIE2-mutant EC (13, 22–24), which demonstrated increased signaling through the PI3K/AKT/mTOR pathway (12, 14, 15).

Unsurprisingly, mutations in the *PIK3CA* gene, which lead to PI3K/AKT hyperactivation, have been identified in approximately 20-30% of sporadic VM cases (25–27), further illustrating the importance of this pathway in promoting VM. Mutant *PIK3CA*-driven VM has been successfully modeled using genetic mouse lines (25, 26, 28–30).

Activation of the PI3K/AKT/mTOR pathway can be targeted by treatment with the mTOR inhibitor rapamycin (20–22, 24, 25), which led to the use of rapamycin in patients with VM and other vascular anomalies (22, 31–34). However, rapamycin has been shown to be incompletely effective in VM patients and in xenograft models of VM (22). While few patients experience partial reductions in lesion volume, rapamycin fails to induce full regression of VM. Furthermore, rapamycin treatment is long-term and cessation often leads to resurgence of symptoms (35).

Investigation of the cellular and molecular mechanisms underlying TIE2-driven venous malformation and the response to therapy has been limited by the lack of a physiologically relevant genetic *in vivo* model. This has also hindered mechanistic studies of disease initiation and progression, as well as preclinical evaluation of targeted therapies. Here, we establish a postnatal endothelial-specific mouse model of TIE2 p.L914F-driven venous malformation that faithfully recapitulates the vascular phenotype observed in patients. Using this model, we define the cellular mechanisms underlying lesion formation and progression and evaluate the therapeutic effects of PI3K/AKT/mTOR pathway inhibition.

## Results

### Systemic postnatal induction of TIE2^L914F^ expression in ECs drives vascular lesion formation

Analysis of cells isolated from sporadic VM patient tissue revealed that the TIE2 p.L914F mutation was found in EC populations and was absent in non-EC types (15). Therefore, to model the sporadic and endothelial origin of VM, we developed a postnatal tamoxifen-inducible murine model expressing TIE2-L914F specifically in EC. We utilized our recently developed mouse line (16), *B6-Tg(Rosa26-TIE2^L914F^)^EBos^*, that contains a Cre-inducible TIE2 p.L914F transgenic construct (**Supplementary Figure S1**), hereafter called *TIE2^L914F^*.

We crossed *TIE2^L914F^* mice to the *Tg(Cdh5-cre/ERT2)^1Rha^* line (hereafter called *iCdh5-Cre*), which expresses a tamoxifen-inducible Cre recombinase specifically in ECs (36) (**Figure 1A**). We then induced transgene expression systemically, through a low dose (0.5µg) intraperitoneal (IP) injection of tamoxifen at postnatal day 5 (P5) (**Figure 1B**).

**Figure 1:**
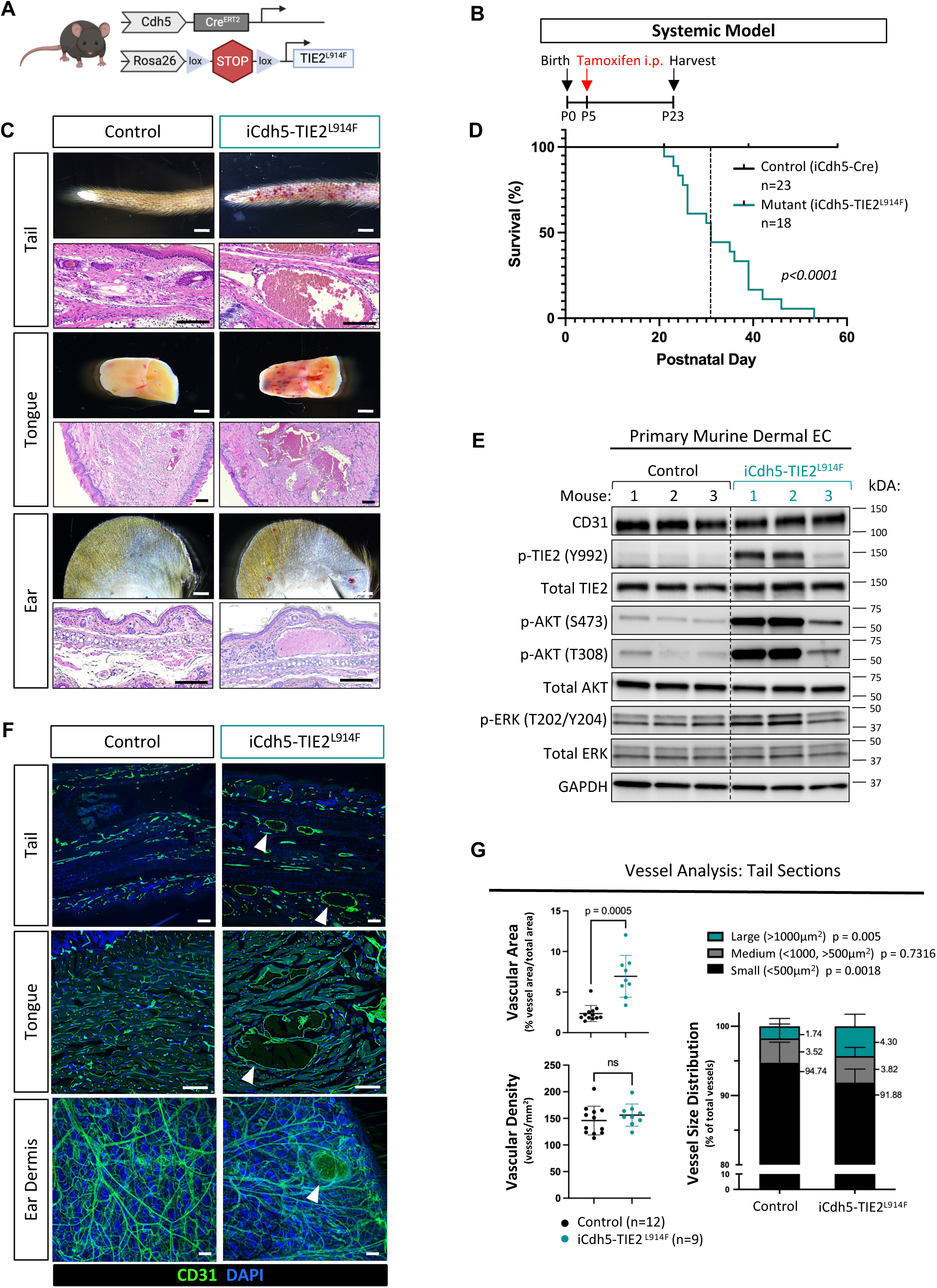
Systemic postnatal induction of TIE2 L914F expression drives formation of widespread vascular lesions. (A) Schematic of the genetic constructs in *iCdh5-TIE2^L914F^*experimental mice. *iCdh5-CreER^T2^* mice were crossed with *TIE2^L914F^* mice to generate *iCdh5-TIE2^L914F^* mutant mice as well as *iCdh5-Cre* controls. (B) Schematic describing the experimental scheme, consisting of tamoxifen induction 5 days after birth (P5) and collection of tissues at P23. (C, top rows) Photographs of gross tissues taken from control and mutant mice at collection at P23. Scale bar = 1mm (white bars). (C, bottom rows) Representative hematoxylin and eosin (H&E) staining of tissue sections, from tissues collected at P23. Scale bar = 100 µm (black bars). (D) Kaplan-Meier curve comparing the survival of *iCdh5-TIE2^L914F^* mutant mice (teal, n=18) and *iCdh5-Cre* control mice (black, n=23). Gehan-Breslow-Wilcoxon test (p < 0.0001). (E) Primary murine dermal EC were isolated from the tail skin of control and mutant mice and cultured *in vitro*. Protein was collected 48h after induction with 4-OHT. Western blot was performed for the indicated phospho- and total proteins. 3 separate mice in each group were used as biological replicates. Nearest molecular weight of ladders are shown to the right. (F) Tail, tongue and ear tissue sections from P23 mice were immunostained for CD31 (green, pan-vascular) and counterstained with DAPI (blue). Scale bar = 100 µm. (G) CD31-stained tail sections were used to quantify vascular parameters. Vascular area is the percentage of vessel area over the total tissue area. Vascular density is the number of vessels per unit of tissue area. Vessel size distribution shows the percentage of total vessels that are considered large (>1000 µm^2^), medium (<1000 µm^2^, >500µm^2^), or small (<500 µm^2^). p-values of the difference between control and mutant mice are indicated for each vessel group in the legend. n=12 control mice and n=9 mutant mice. All data are shown as mean±SD, Welch’s t-test and two-way Anova for vessel size distribution analysis. Schematic in (A) created with Biorender.com.

We found that these mice developed multiple vascular lesions that preferentially appeared in skin and muscle tissue, such as those of the tail, tongue, and ear (**Figure 1C, top rows**). Histology of those tissues revealed the presence of abnormal vessels with massively enlarged lumens (**Figure 1C, bottom rows**). In addition, we found that the widespread and pervasive nature of lesions in mutant mice limited survival, with the mutant *iCdh5-TIE2^L914F^* mice having 50% survival at P30 (**Figure 1D**).

To confirm activation of TIE2 signaling, we isolated dermal EC from *iCdh5-TIE2^L914F^* and control mice. Cells were cultured *in vitro* and transgene expression was induced with 4-hydroxytamoxifen (4-OHT). We then performed Western blotting analysis to investigate phosphorylation of TIE2 and its downstream signaling proteins. EC from iCdh5-*TIE2^L914F^* mice showed increased phosphorylation of TIE2, AKT, and ERK compared with controls (**Figure 1E**).

Having established that *iCdh5-TIE2^L914F^* mice develop a vascular phenotype that is associated with increased TIE2 signaling in EC, we then sought to evaluate the vascular changes happening in affected tissues. We performed immunostaining of tail, tongue and ear tissues for CD31, a marker specifically expressed by EC and show enlarged and dysmorphic vessels in mutant mice compared to controls (**Figure 1F**). Quantification of the CD31^+^ vasculature in tail sections revealed that the vascular area was increased without an increase in vessel density. Furthermore, analysis of vessel size distribution showed an increase in the number of large (>1000μm^2^) vessels, with a corresponding decrease in the number of small vessels (<500μm^2^) (**Figure 1G**). This result is consistent with enlargement of pre-existing vessels rather than increased formation of new vessels.

In total, this model demonstrates that postnatal expression of TIE2 p.L914F in EC is sufficient to drive formation of a VM-like phenotype, with dilation of blood vessels that contributes to an increase in vascular area, but not an increase in vascular density. Furthermore, EC from *iCdh5-TIE2^L914F^* mice show increased TIE2 activation and downstream signaling, confirming activation of the expected downstream signaling pathways *in vivo*.

### Robust formation of visible VM lesions within ear skin is driven by topical induction of TIE2^L914F^ in ECs

While we gathered significant information from the systemic tamoxifen model, we encountered phenotypic variability. While lesions consistently appeared in mutant mice, the location and timing of these lesions was variable between individual mice, prohibiting time-resolved studies of the cellular changes associated with lesion formation and progression. To better mimic the somatic nature of VM, we induced Cre activation by topically applying active tamoxifen (4-OHT) directly to the ears of P10 mice (**Figure 2A**). This approach drove robust and consistent vascular lesion formation in the ear skin of mutant mice (**Figure 2B**).

**Figure 2:**
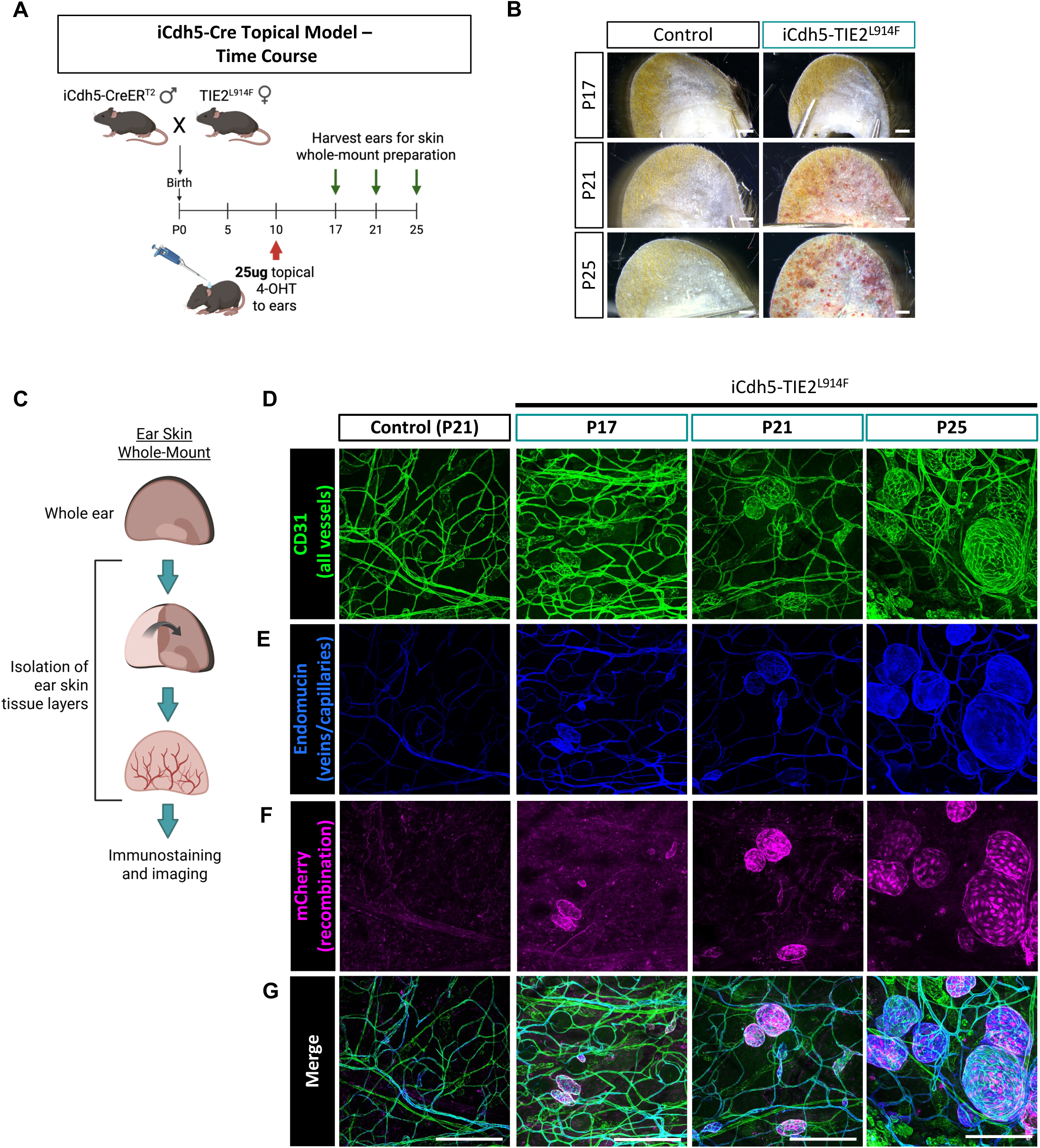
Topical administration of 4-OHT to ears of postnatal iCdh5-TIE2^L914F^ mice drives mosaic recombination and lesion formation within the venous/capillary plexus. (A) Schematic describing the mouse model for time course analysis. *iCdh5-CreER^T2^*mice were crossed with *TIE2^L914F^* mice and offspring were induced with 25µg 4-OHT via topical administration to the ears at P10. The ears of mice were then collected at P17, P23, and P25. (B) Representative photographs of ear tissues collected from control and mutant mice at each time point. Scale bar = 1mm. Representative of n=4-8 per group. (C) Schematic showing the isolation of ear skin whole-mounts from whole ear tissue. (D-G) Representative images of ear skin whole-mounts from each time point that were immunostained for CD31 (green, pan-vascular), Endomucin (blue, veins/capillaries), mCherry (magenta, recombined cells) and merge. Scale bar = 250 µm. Schematics in (A and C) created with Biorender.com.

Based on the timing of visible lesion formation in the ears, we performed tissue collection at P17, P21, and P25, timepoints that encompass early lesion formation (P17) and lesion expansion (P21 and P25) (**Figure 2B**). To visualize the vasculature, we performed immunostaining and confocal imaging of skin whole-mounts from the ears (**Figure 2C**). CD31 staining revealed abnormal cyst-like focal enlargements of the blood vessels in the *iCdh5-TIE2^L914F^* mice as early as P17 with progressive enlargement of lesions through P21 and P25 (**Figure 2D**). To determine whether the lesional vessels were of venous origin, recapitulating the venous specificity seen in VM patients, tissues were stained for Endomucin, which we confirmed as a marker of venous/capillary EC in wild-type murine tissue (37) (**Supplementary Figure S2**). Ear skin VM lesions were universally positive for Endomucin, (**Figure 2E**), illustrating that the mutation is preferentially driving lesion formation in the venous/capillary plexus. Similar results were obtained in the systemic model, where enlarged lesional vessels in ear, tongue, and tail stained positively for Endomucin (**Supplementary Figure S3**).

To determine unequivocally that lesions were formed of *TIE2^L914F^* mutant EC, we stained the tissue for mCherry (**Figure 2F**), a reporter included in our transgenic construct (**Supplementary Figure S1**). *iCdh5-TIE2^L914F^* lesions were overwhelmingly composed of mCherry^+^ EC, indicating that the lesions are a direct consequence of TIE2-mutant EC. Furthermore, the mCherry^+^ EC were localized within the lesional Endomucin+ capillary/venous vessels (**Figure 2G**). These results suggest that clonal expansion of the mutant EC drives VM lesion formation. To confirm that this finding also applies to the systemic model, we examined tongue sections from control and mutant mice treated with systemic tamoxifen. Here, we also found that lesional vessels were lined by mCherry^+^ (TIE2-mutant) EC (**Supplementary Figure S4**), indicating that lesions are formed by mutant EC with a similar mechanism of clonal expansion, regardless of the route by which tamoxifen is administered.

With the topical 4-OHT administration, our aim was to induce local lesion formation circumventing the deleterious effects of systemic tamoxifen delivery. which induces widespread lesion formation and 50% lethality around P30 (as seen in **Figure 1B, D**). However, we found that mutant mice administered topical 4-OHT still developed lesions throughout the body, leading to eventual moribundity and death, with median survival at P27 (**Supplementary Figure S5**). One possible explanation is that 4-OHT enters the systemic administration, as has been reported in other topical models (38).

These results demonstrate that topical administration of tamoxifen onto a specific tissue drives more robust and reproducible formation of vascular lesions, allowing for detailed and time-resolved analysis of phenotypes. Furthermore, the appearance of visible lesions on the ear allows for longitudinal imaging of lesion development in live mice.

### Expression of TIE2^L914F^ in veins, but not arteries or lymphatics, is sufficient to drive vascular lesion formation

Our data indicate that TIE2^L914F^-driven lesions preferentially form in the Endomucin^+^ venous/capillary plexus, despite the *iCdh5-Cre* driver being a pan-EC Cre driver (36). To determine whether venous-specific expression of the mutation is sufficient to induce lesion formation we crossed the *TIE2^L914F^*mice with the *Gm5127(BAC)CreER^T2^* line (hereafter called *iV(enous)EC-Cre*), which has been shown to be specific for venous endothelial cells (VEC) of the dermis, brain, and retina (39). In this model, we induced recombination with topical application of 4-OHT to the ears of control and mutant *iVEC-Cre; TIE2^L914F^* mice (**Figure 3A**). Mutant mice developed visible vascular lesions on the ears by P52 (**Figure 3B**). Histology of ear sections further showed the presence of Endomucin^+^ vessels with massively enlarged lumens in *iVEC-Cre; TIE2^L914F^* mice (**Figure 3C**), resembling those found in *iCdh5-TIE2^L914F^* mice. The percent lesion area, calculated from photographs of collected ears, showed that lesions formed in the majority of *iVEC-Cre; TIE2^L914F^* mice, with no lesions forming in Cre+ control mice that also received 4-OHT (**Figure 3D**).

**Figure 3:**
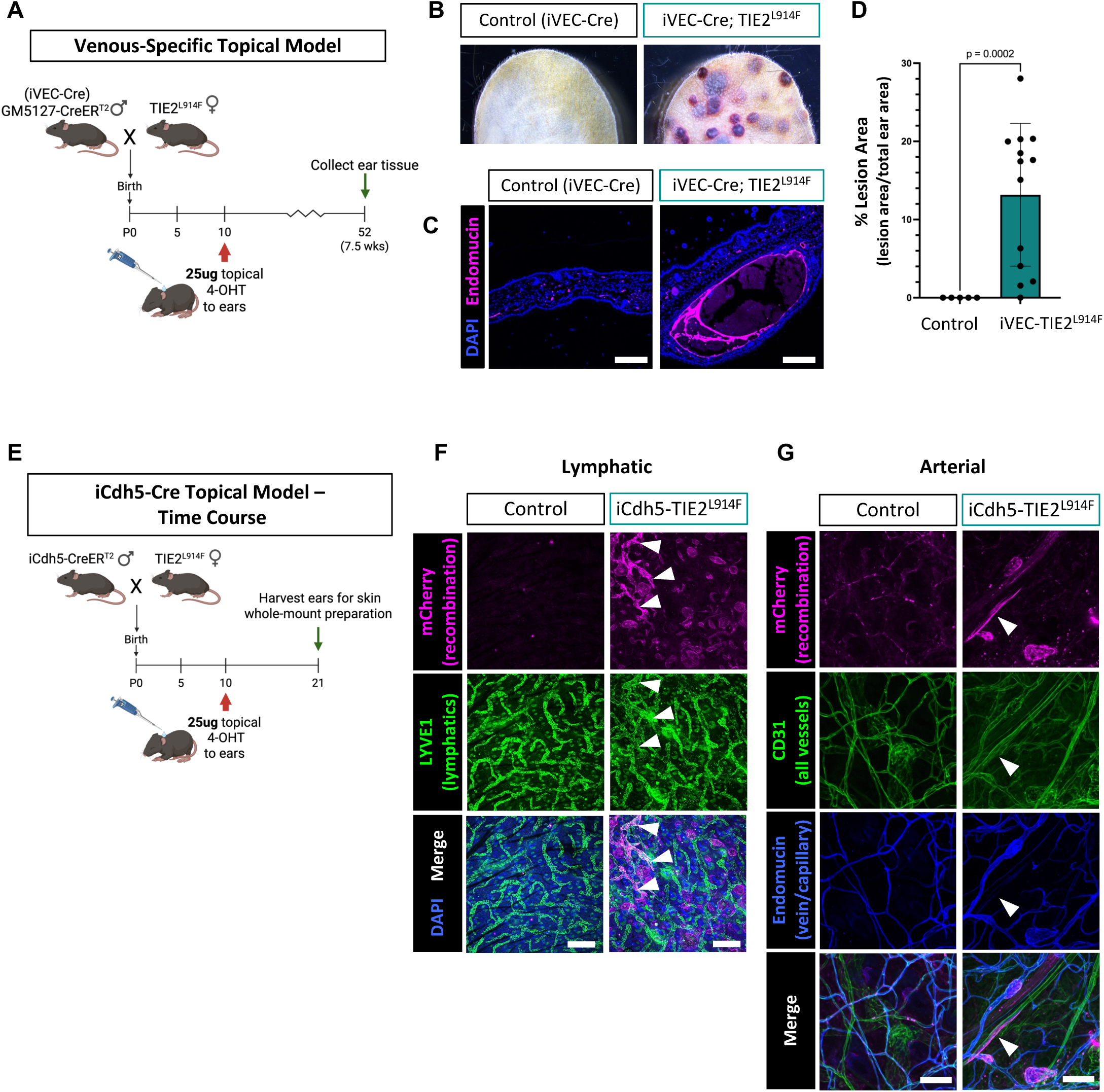
Venous-specific TIE2 L914F expression is sufficient to form vascular lesions while TIE2 L914F-expressing arteries and lymphatics fail to form lesions. (A) Schematic describing the generation of a venous-specific mouse model. *iVEC-Cre* mice were crossed with *TIE2^L914F^* mice and offspring were induced with 25µg 4-OHT via topical administration to the ears at P10. Ear tissue was then collected at P52. (B) Representative photographs of ear tissue taken from control (*iVEC-Cre*) and mutant (*iVEC-TIE2^L914F^*) mice. (C) Ear tissue sections were immunostained for Endomucin (magenta, veins/capillaries) and counterstained with DAPI (blue, nuclei). Scale bar = 250 µm. (D) The percent lesion area was measured from photographs of mouse ears and is shown as the total lesional area over the total area of the ear. Mean±SD, Welch’s t-test. n=5 control mice and n=13 mutant mice. (E) Schematic describing the *iCdh5-TIE2^L914F^* experimental topical model. Offspring were induced with 25µg 4-OHT via topical administration to the ears at P10 and tissues were harvested at P21. (F) Ear skin whole-mounts from control and mutant mice were immunostained for mCherry (magenta, recombined cells), LYVE1 (green, lymphatic vessels), and counterstained with DAPI (blue, nuclei). Representative of n=3 mice. Scale bar = 250 µm. (G) Ear skin whole-mounts from control and mutant mice were immunostained for mCherry (magenta, recombined cells), CD31 (green, pan-vascular), and Endomucin (blue, veins/capillaries). Arteries were defined as CD31^+^ but Endomucin^-^. Representative of n=5 mice. Scale bar = 250 µm. Schematic in (A and E) created with Biorender.com.

Having shown that venous-restricted expression of mutant TIE2 is sufficient to drive lesion formation, we next asked whether expression in lymphatic or arterial EC could also alter vessel morphology. We therefore returned to the pan-EC *iCdh5-Cre* topical model, at the P21 timepoint, and analyzed the ear skin whole-mounts for the presence of non-venous/capillary mCherry^+^ EC (**Figure 3E**). We first co-stained the tissue for LYVE1, a marker of lymphatics, as well as mCherry. Within *iCdh5-TIE2^L914F^* mouse ears, we identified mCherry^+^ lymphatics that were non-lesional and similar in structure and size to wild-type (WT) mCherry^-^ lymphatics (**Figure 3F**). Next, we co-stained for Endomucin, CD31, and mCherry and analyzed the tissue for mCherry^+^ arterial (CD31^+^ Endomucin^-^) ECs. These arterial vessels were also non-lesional and similar to mCherry^-^ (WT) arteries (**Figure 3G**).

One proposed explanation for the venous specificity of TIE2-p.L914F-driven lesions is that venous EC are intrinsically more proliferative than arterial EC (39). To test this hypothesis, we quantified baseline EC proliferation in venous, arterial, and lymphatic vessels. Control mice at P17 (the timepoint at which lesions begin to form in mutant mice) were administered EdU 24h before tissue collection to label proliferating cells. Their ear skin whole-mounts were stained for markers of venous (Endomucin), arterial (SOX17), and lymphatic (PROX1) EC (**Supplementary Figure S6**). Quantification of the number of proliferating EC in each vessel type showed that, at this timepoint, the ear skin lymphatic EC were the most highly proliferative, followed by venous/capillary EC, and finally arterial EC. Therefore, while the low proliferative capacity of arterial ECs may explain why they are not found in *TIE2^L914F^*-driven lesions, the high proliferative capacity of lymphatic vessels suggests additional mechanisms can cause LEC to be resistant to *TIE2^L914F^*-driven lesion formation.

Together, these findings demonstrate that TIE2 p.L914F preferentially induces lesions in venous EC and indicate that differences in baseline EC proliferation alone are insufficient to explain this vessel-type specificity.

### Venous malformation lesion formation in *iCdh5-TIE2^L914F^* mice is associated with a biphasic increase of EC proliferation and cell enlargement

Having established the vessel-of-origin of the TIE2-driven vascular lesions, we then sought to determine the cellular processes that could be driving vessel enlargement. To investigate cell proliferation, we used the previously established time course model (**Figure 4A**), and we injected EdU into mice 24h before each collection. We then performed immunostaining of ear skin whole-mounts for Endomucin (veins/capillaries), ERG (EC nuclei), and EdU (proliferating nuclei). To control for baseline differences in vessel type, we masked the nuclear signals on the Endomucin signal, so that we would only measure proliferating EC in veins and capillaries (**Figure 4B**). Here we observed an increase in the number of triple-positive proliferating EC nuclei (EdU^+^ ERG^+^ Endomucin^+^) in *iCdh5-TIE2^L914F^* mice at each timepoint, many of which were concentrated within the lesional vessels. We further quantified the number of triple-positive nuclei as a percentage of total Endomucin^+^ ERG^+^ (venous/capillary EC) nuclei and found that EC proliferation was increased in the mutant mice at each timepoint (**Figure 4C**). However, the magnitude of the increase in proliferation in the mutant mice compared to controls decreased at the latter two timepoints (P21 and P25) compared to the first timepoint (P17). At P17, the excess cell proliferation that can be attributed to the mutation is 10% (15% proliferating ECs in mutant tissue minus 5% baseline proliferation in control tissues). However, the excess cell proliferation at P21 and P25 is only about 5-6%, despite the fact that there are many more lesional EC at these later timepoints.

**Figure 4:**
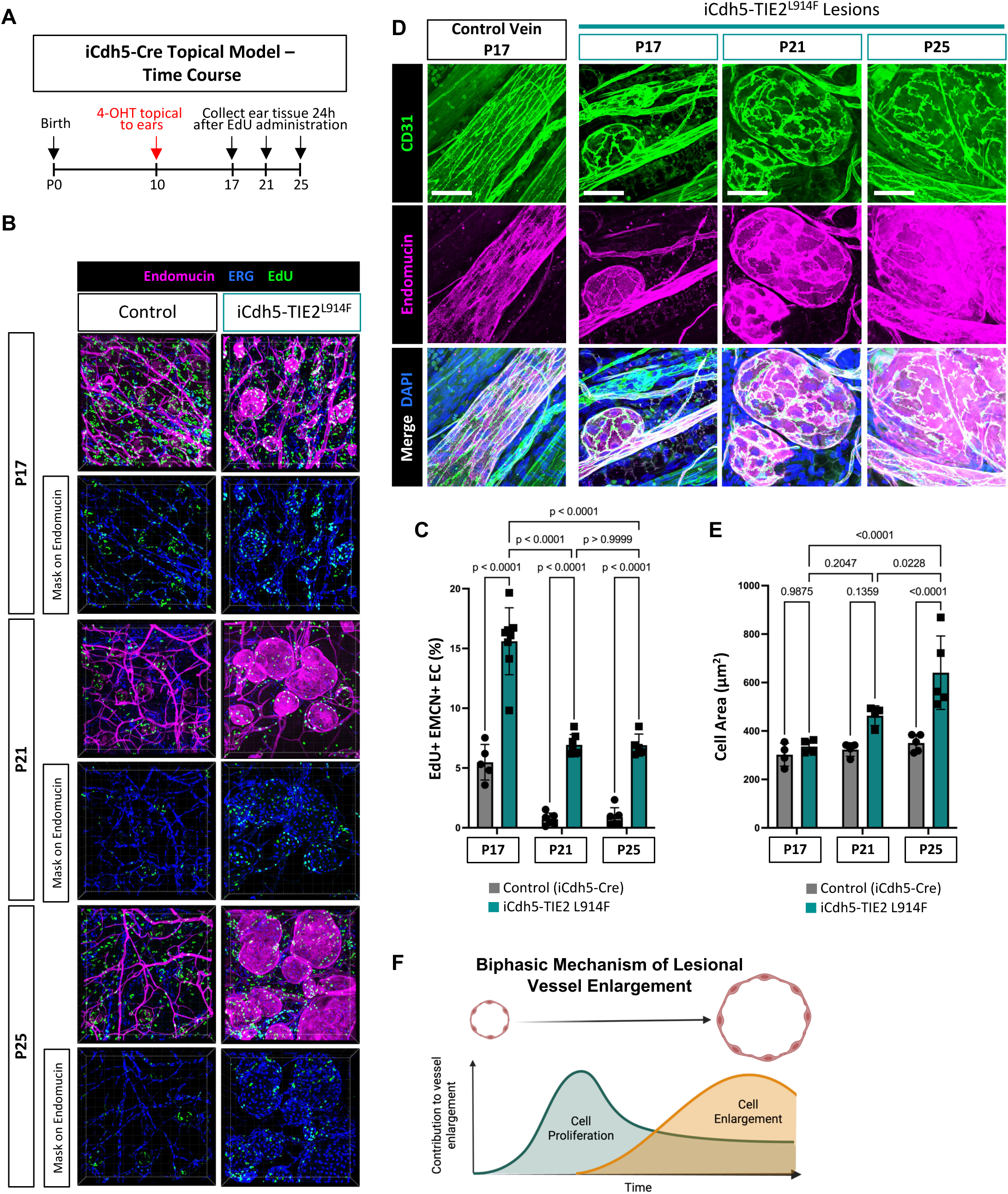
Lesion formation and progression are associated with a biphasic increase in EC proliferation and cell size. (A) Schematic showing experimental protocol. Mice were administered 25µg of 4-OHT topically to the ears at P10. Tissues were collected at P17, P21, and P25. EdU was injected into mice 24h before each collection to label proliferating cells. (B) Representative immunofluorescent images of ear skin whole-mounts. Veins/capillaries are labeled with Endomucin (magenta), EC nuclei with ERG (blue), and proliferating cells with EdU (green). Masking of nuclear signals on Endomucin channel was performed using Imaris software. (C) The number of EdU+ EMCN+ ERG+ nuclei was quantified using General Analysis 3 in Nikon Elements Software and is represented as a percentage of total EMCN+ ERG+ nuclei. n=5-8 mice per group. Mean±SD. Two-way ANOVA with multiple comparisons. (D) Ear whole mounts were also immunostained for Endomucin (magenta, veins/capillaries) and CD31 (green, EC cell borders). Scale bar = 50μm. (E) CD31+ cell borders were manually outlined to quantify cell area of non-lesional and lesional vessels in *iCdh5-TIE2^L914F^* mice. Average cell area for individual mice is shown, n=4-5 mice per group. Mean±SD. Two-way ANOVA with multiple comparisons. (F) Schematic showing biphasic contributions of EC proliferation and enlargement to lesional vessel enlargement. Created with Biorender.com.

Because of this reduction in the lesional EC proliferation rate, we sought to determine whether additional cellular processes could contribute to VM expansion. We hypothesized that changes in EC size could play a role in VM lesion growth, as it has been shown to be increased in other animals models of vascular anomalies (including hereditary hemorrhagic telangiectasia and arteriovenous malformation) (40–42), as well as in an *in vitro* model of TIE2^L914F^ EC under flow(14). We used CD31^+^ cell borders to measure EC area as a proxy for EC size in normal veins and in lesional vessels (**Figure 4D**). Quantification of cell area revealed that while lesional EC were similar in size to non-lesional EC at the P17 timepoint, the lesional EC became progressively larger through the P21 timepoint, and were significantly larger (p<0.0001) by the P25 timepoint (**Figure 4E**).

In total, this analysis of both cell proliferation and cell size in a time course model that encompasses both early, middle, and late lesion formation reveals a biphasic contribution to lesion enlargement (**Figure 4F**). In TIE2 p.L914F-mutant mice, EC proliferation in lesional vessels is highest at the earliest stages of lesion formation, when lesions are the smallest in size (P17). However, at this same timepoint, EC size is not significantly changed. While a mildly elevated level of proliferation is maintained at later timepoints (P21 and P25), the magnitude of the increase was not as large as at P17. This is especially striking considering that there are many more lesional EC present at the later stages due to lesion expansion, which suggests that the overall percentage of mutant cells that are actively proliferating is even more reduced at later stages. However, EC size does become progressively larger throughout the later time points.

Our results show a biphasic mechanism is responsible for the pathogenic vessel enlargement. This indicates that different abnormal cellular phenotypes contribute to lesion enlargement at different points in the process of lesion formation and progression.

### *TIE2^L914F^*-driven vascular lesions are associated with irregular vascular smooth muscle cell coverage

Histopathology of patient VM lesions revealed that the abnormal veins often exhibit changes to the surrounding mural cells, including disorganized and uneven coverage by the vascular smooth muscle cells (vSMC) that would typically support a large vein (8, 9, 43, 44). To examine whether vSMC coverage of lesional vessels was affected in our model, we stained ear skin whole-mounts from the *iCdh5-Cre* topical time course model (**Figure 5A**) for αSMA (smooth muscle alpha actin), a marker of vSMC. We first looked at P17, when lesions are in the early stages of formation, and found that small lesional vessels had inconsistent coverage of vSMC, even when budding off parental veins with high and organized coverage of vSMC (**Figure 5B**). We then examined vSMC coverage at P21, when lesions expanded in size, and found further reduced vSMC coverage of lesions, which when quantified showed a significant (p=0.0253) decrease compared to control veins (**Figure 5C**).

**Figure 5:**
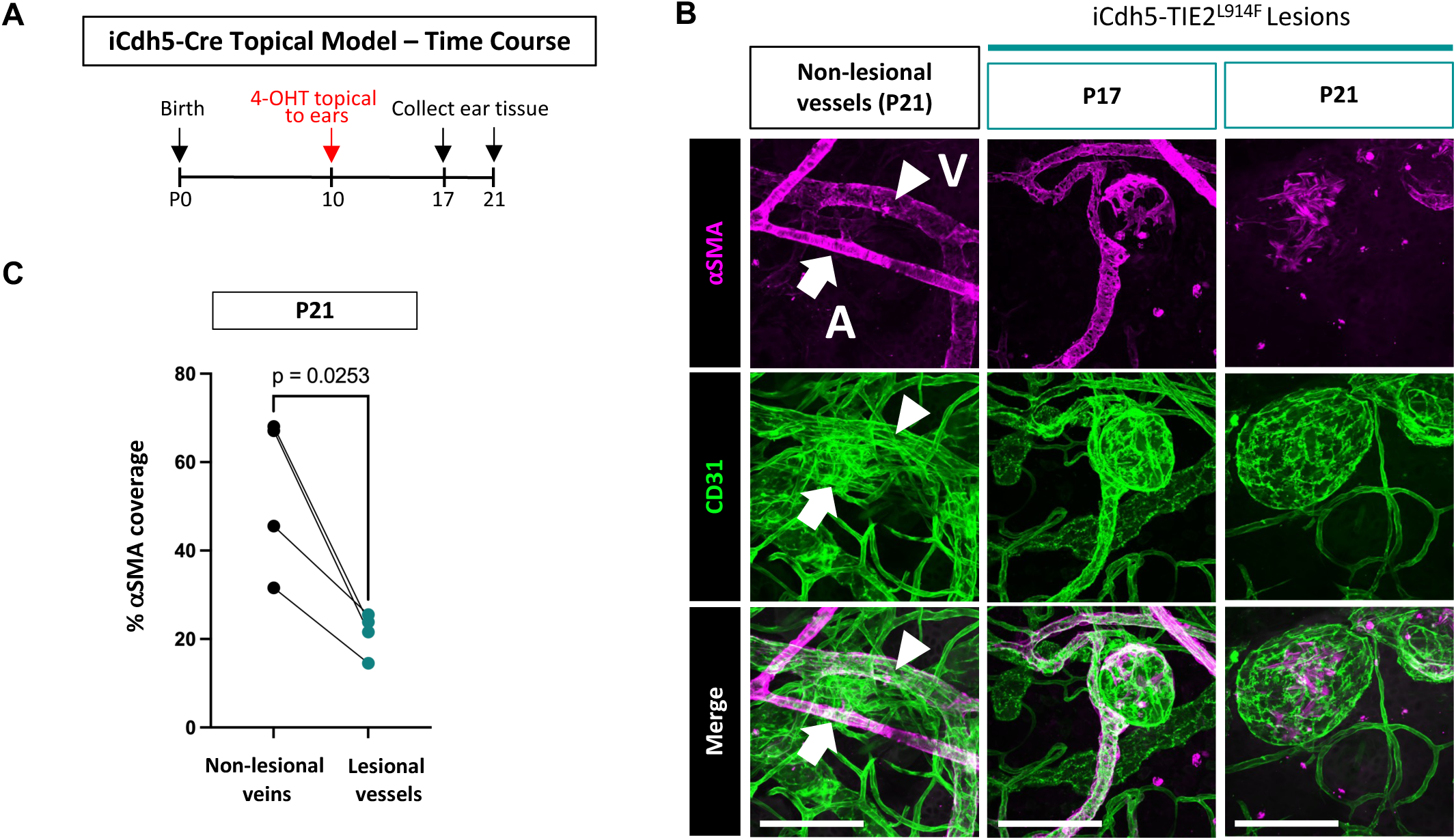
Abnormal vessels in iCdh5-TIE2^L914F^ mice have reduced coverage of smooth muscle cells. (A) Schematic of experimental protocol. The time course scheme was used, in which mice were treated with 25µg of 4-OHT topically to the ears at P10 and tissues were collected at P17 and P21. (B) Representative images of ear skin whole-mounts taken from P17 and P21 mice immunostained for CD31 (pan-vascular, green) and *a*SMA (vSMCs, magenta). Scale bars = 100μm. (C) The percent of *a*SMA coverage on non-lesional and lesional vessels within the same mouse (connected by lines) was analyzed from the immunostaining in panel B. n=4 mice, 5-18 vessels per group per mouse. Mean±SD. Paired t-test.

Together, these findings indicate that TIE2-L914F-driven VM lesions are associated with progressive disruption of vSMC coverage. This irregular mural cell investment during lesion progression phenocopies patients’ VM lesions histopathology.

### Rapamycin treatment reduces lesion growth by partially rescuing abnormal EC proliferation in *iCdh5-TIE2^L914F^* mice

TIE2 p.L914F has previously been shown to increase TIE2 signaling through the PI3K/AKT/mTOR pathway (12, 45) which can be successfully downregulated by treatment with the mTOR inhibitor rapamycin (22, 46). These discoveries led to the use of rapamycin in the treatment of VM patients with some success, in that patients report improved quality of life and moderate reduction in lesion volume. However, full lesion regression is typically not achieved (22, 34).

Therefore, we sought to determine the phenotypic and cellular effects targeted by rapamycin treatment in the *iCdh5-TIE2^L914F^* model. First, we confirmed that rapamycin treatment *in vitro* reduces the increased levels of phospho-AKT in dermal EC isolated from *iCdh5-TIE2^L914F^*mice (**Supplementary Figure S7A-B**). Having shown that rapamycin is effective at inhibiting AKT signaling in EC, we then established an *in vivo* rapamycin treatment scheme. We administered 4-OHT to the ears of mice at P10, but at a lower dose (5μg), which allowed for longer survival and for chronic drug treatment regimens, more closely mimicking the clinical scenario. We then administered 2mg/kg of rapamycin every other day starting at P12 and throughout the course of the experiment (**Figure 6A**). Because the lesions are visible on the ears of mice, we took photographs twice a week, allowing us to monitor lesion growth longitudinally.

**Figure 6:**
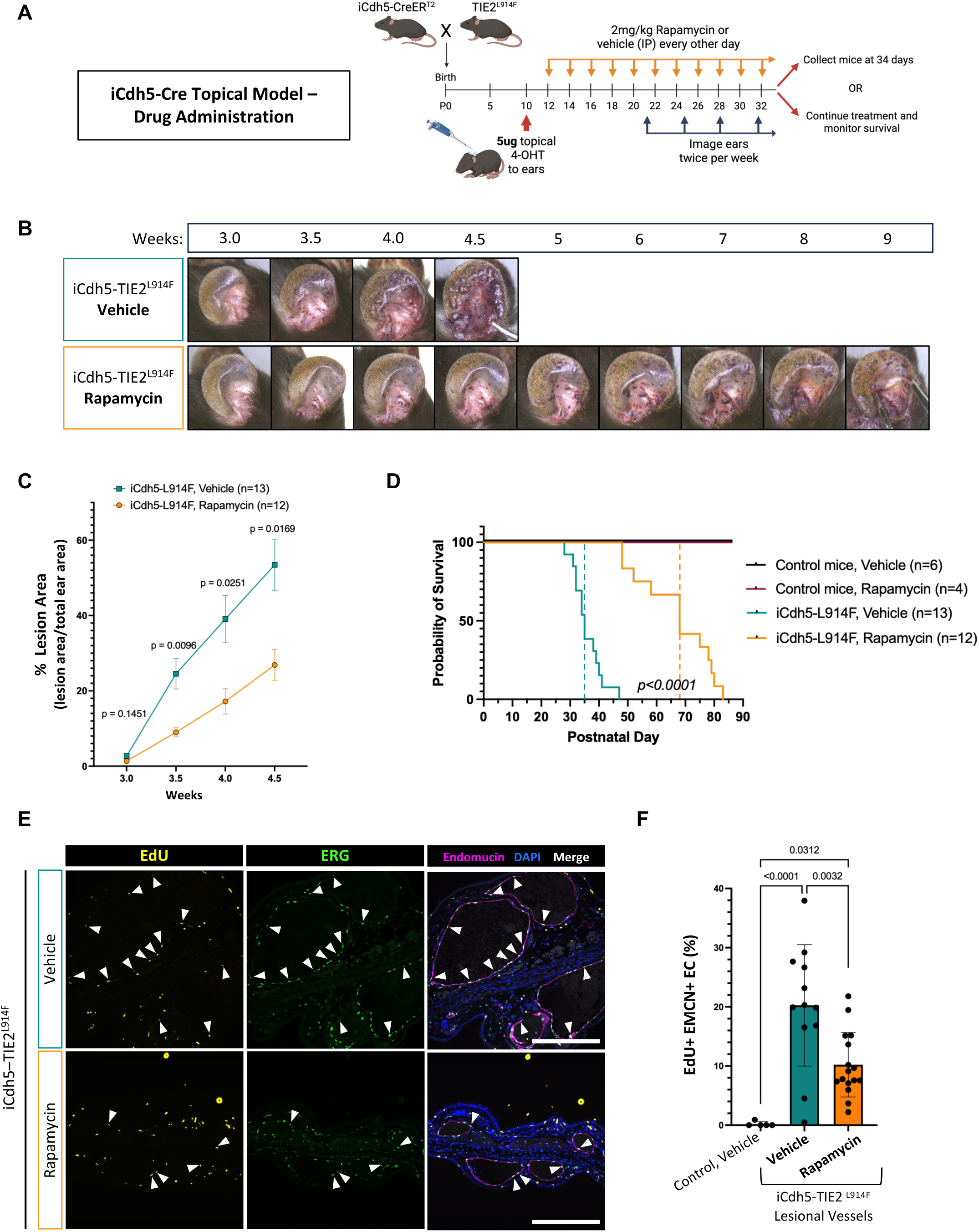
Increased activation of PI3K/AKT/mTOR signaling in iCdh5-TIE2^L914F^ mice can be targeted with rapamycin to reduce lesion growth, extend survival, and reduce EC proliferation. (A) Schematic of experimental setup. Mice were administered 5μg of 4-OHT topically to the ears at P10. Rapamycin (2mg/kg, i.p,) was administered every other day starting at P12. The ears of live mice were imaged twice per week starting at 3 weeks of age (P21). (B) Images of ears were obtained on live mice using a stereomicroscope. Individual mice were imaged longitudinally over time. (C) The percentage of lesion area over total ear area was quantified from images in panel C. Mean±SEM. Mixed-effects analysis with Sidak’s multiple comparisons test. (D) Kaplan-Meier curve comparing the survival of vehicle- and rapamycin-treated *iCdh5-TIE2^L914F^* mutant and *iCdh5-Cre* control mice. Number of mice for each group is indicated in the figure legend. Gehan-Breslow-Wilcoxon test (p<0.0001). (E) Sections of ear tissue were stained for Endomucin (magenta, veins/capillaries), ERG (green, EC nuclei), and EdU (yellow, proliferating cells). Endomucin+ ERG+ EdU+ cells within lesions are indicated by arrows. Scale bar = 250μm. (F) The number of EdU+ EMCN+ ERG+ nuclei within lesions was quantified and is represented as a percentage of total lesional EMCN+ ERG+ nuclei. Vessels over 30 μm in diameter were considered lesional. n=5-16 mice per group. Mean±SD. One-way ANOVA with multiple comparisons. Schematic in (A) created with Biorender.com.

Vehicle-treated mutant mice developed lesions that appeared around 3 weeks of age and grew rapidly, eventually encompassing most of the ear (**Figure 6B-C**) and leading to mortality from P30-50, with a median survival occurring at P35 (**Figure 6D**). Rapamycin-treated mutant mice also developed visible vascular lesions (**Figure 6B**), but these grew at a significantly reduced rate (**Figure 6C**), prolonging mouse survival up to P80, with median survival occurring at P68 (**Figure 6D**). We did not observe changes in survival or weight in response to rapamycin treatment attributable to male or female sex (**Supplementary Figure S8**). These data establish our genetic model of VM as an important tool for preclinical studies. In this model, lesions can be analyzed longitudinally and we observed a response to rapamycin similar to VM patients.

We next examined the effects of rapamycin on the two cellular processes underlying lesion formation and expansion: endothelial cell proliferation and cell size. To do so, we collected ear tissue from control and treated mice at P34, before there is high (>50%) mortality rate in the vehicle-treated group (**Figure 6D**). We found a significant (p=0.0032), but partial, reduction in the proliferation of lesional EC in rapamycin-treated *iCdh5-TIE2^L914F^* mice (**Figure 6E-F**). However, we found no difference in the size of lesional EC in rapamycin-treated mice compared to vehicle-treated (**Supplementary Figure S9**). Furthermore, we observed that lesions from rapamycin-treated mice were still deficient in vSMC coverage (**Supplementary Figure S10**). Finally, we found that rapamycin-treated mice exhibited mild but consistent reductions in weight gain, which has been observed by other groups utilizing early-life treatment of rapamycin to mice(47–49) (**Supplementary Figure S8B-C**), but was otherwise well tolerated.

Together, these findings demonstrate that rapamycin partially suppresses TIE2-L914F-driven lesion progression by reducing EC proliferation, while failing to rescue EC hypertrophy.

## Discussion

Here we report a postnatal endothelial-specific TIE2-L914F model that not only recapitulates human VMs but also provides novel insights into the mechanisms of lesion development. Using this model, we show that TIE2-driven lesions arise selectively within the venous vasculature and undergo clonal expansion. Dysmorphic vessels originate from a biphasic program of endothelial cell proliferation followed by cell enlargement and exhibit progressive mural cell abnormalities. Therapeutically, we show that TIE2-L914F venous malformations respond PI3K/AKT/mTOR inhibition, similarly to patients.

An ongoing area of investigation in the vascular anomalies field is the biological basis for the vessel-type specificity that is seen with various mutations. For VM specifically, genetic studies in patients have revealed the presence of TIE2 p.L914F mutations only in VM, whereas other mutations occur in multiple types of vascular malformation, such as *PIK3CA* mutations that have been found in VM, lymphatic malformations, and other types of malformations and overgrowth syndromes (50). However, the specificity of TIE2 p.L914F for venous lesions has never been shown in experimental systems, precluding investigations of its vessel type-specific effects. Therefore, in this study, using a venous-specific Cre-driver mouse line (*iVEC-Cre*), we show that expression of TIE2 p.L914F in veins is sufficient to drive vascular lesion formation. Perhaps even more interestingly, in *Cdh5-TIE2^L914F^* mice, where recombination occurs in all types of EC, we determined that transgene (mCherry) expressing arterial and lymphatic vessels were morphologically normal. In patient VM, it could be postulated that lack of expression of the mutation in these vessel types prevents lesion formation (such as from lower expression of the *TEK* gene or mutation-induced cell death). However, our data does not support this hypothesis. Rather, it appears that the EC in arteries and lymphatic vessels are inherently resistant to the effects of TIE2 p.L914F mutation. Even upon forced expression of the mutant gene in these vessels, they fail to experience the same lumen expansion that occurs when the mutation is expressed in veins/capillaries. One possible hypothesis to explain this difference is the inherent proliferative capacity of the various vessel types, as it has been shown that venous EC are typically more proliferative (at least than arterial) and are often the source of angiogenic EC in physiological and pathological vascular processes (39). However, the results obtained in the ear vasculature with our model do not support this hypothesis. We found that in the ear skin, at the timepoints used in our experiments, lymphatic EC are the most proliferative and there are still detectable amounts of proliferating arterial EC. Therefore, we think it is likely that additional factors such as cell identity, flow mechanics, expression of regulators of TIE2 signaling, and other factors are more likely to be mediating this effect. Some hints are offered by *PIK3CA* mutant models, where recent work has shown that mature arterial cells of the retina are inherently refractory to the effects of *PIK3CA^H1047R^*mutation and that its expression in pre-arterial cells drives switching to a venous/capillary fate (51). Future work in the *TIE2^L914F^* mouse model and in other systems will be needed to determine if these mechanisms and/or others are involved.

Using the *Cdh5-TIE2^L914F^* model, we also investigated the cellular changes that occur during lesion formation and progression. Based on mCherry reporter activity that shows where expression of the transgenic construct is occurring, we observed clonal expansion of lesions within existing vessels, which has also been shown in a xenograft model with TIE2 mutant EC (23) and in PIK3CA-driven vascular malformation models (30). This is supported by the observation that there are very high and abnormal levels of EC proliferation within the TIE2 mutant vascular lesions. However, we did also observe that EC proliferation in mutant vessels was highest at the earliest timepoint of lesion formation (P17), despite there being fewer lesional EC. At the later timepoints (P21 and P25) the pool of mutant EC is greatly expanded, but fewer of those cells appear to be proliferating. Therefore, these data suggest that there is an initial burst of EC proliferation that occurs during early lesion formation, that is then sustained at lower levels throughout lesion progression. Clinically, lesions are surgically resected only when they reach a certain size and are symptomatic and this may explain why historically, when compared to vascular tumors, EC proliferation in VM has not been considered pathological. We speculate that as lesions increase in size, proliferation slows down and changes to vessel mechanics or to blood flow-induced shear forces may have impacts on cell signaling and phenotype. Furthermore, we found that the increased EC proliferation in VM lesions was partially rescued by rapamycin, implicating PI3K/AKT/mTOR signaling in promoting this phenotype. This would suggest that increased treatment efficacy is associated with early administration of rapamycin, when lesions are still small and developing.

We also observed an increase in cell size during VM lesion expansion, with lesional EC becoming progressively larger than normal venous EC throughout lesion growth, although they are not significantly larger at the initial timepoint of lesion formation (P17). This suggests that EC size is not likely a significant contributing factor during lesion formation (perhaps due to the high rates of EC proliferation preventing excessive cell enlargement) but at later stages may contribute to lesion expansion. While it is possible that the EC enlargement may be a consequence of lesion formation rather than a cause, we find that there is additional evidence from the literature to support its potential role in lesion growth. Other models of vascular anomalies have shown changes to EC size that have been involved in formation of abnormal vessels (40–42) and there is *in vitro* evidence suggesting that TIE2^L914F^-expressing EC autonomously increase in cell size in a microfluidic model with flow-induced shear stress (14). Interestingly, we did not see a rescue of the abnormal EC size upon rapamycin treatment, despite lesions being smaller. This data suggests that the increase in EC size is not entirely dependent on the size of the lesion and that, if driven by cellular signaling, it is not dependent on the PI3K/AKT/mTOR pathway. Therefore, future work will be needed to determine whether changes to EC size in our model may be dependent on other signaling pathways or may be caused by the aberrant blood flow and shear stress that occurs in lesions, regardless of their size.

In conclusion, we established a novel genetic murine model that recapitulates TIE2 p.L914F-driven VM disease in mouse. We show that it models key characteristics of patient disease, including formation of visible vascularized lesions, pathological vessel enlargement in relevant tissues, venous specificity, and abnormal recruitment of vSMC. We also identify cellular mechanisms of lumen expansion, including a biphasic increase in EC proliferation and subsequent cell size enlargement. Finally, we determined that rapamycin is capable of partially rescuing vascular lesion through its ability to reduce abnormal EC proliferation, but not EC enlargement. While further work is needed to find additional, targetable signaling pathways that drive the phenotypes not affected by rapamycin treatment, this model will provide a key experimental tool in the investigation of those pathways and will provide an ideal platform for testing of new therapies for the treatment of VM.

## Methods

### Mouse husbandry and experimental procedures

#### Genotyping and mouse maintenance

B6-Tg(Rosa26-TIE2^L914F^)^EBos^ (*TIE2^L914F^*) mice were generated as described previously (16). For routine genotyping of colony and experimental mice, PCR primers were used as indicated in **Table 1** with EconoTaq Plus green 2X Master Mix (Lucigen, cat# 30033). For colony genotyping, murine DNA was extracted from ear biopsies using the protocol established in Truett, et al. (2000)(52). For genotyping of postnatal experimental mice, toe or tail clips from P2-P5 mice were processed using the same protocol.

Mice were housed in the animal care facility of Cincinnati Children’s Hospital Medical Center under standard pathogen-free conditions with a 14 h light/10 h dark schedule and provided with food and water ad libitum. All mice were maintained on a C57BL/6 background and both male and female mice were used in all experiments. All experimental mice were monitored every other day for signs of moribundity, including lethargy, weight loss, and functional impairment from lesion formation. Upon experiment endpoint or moribundity, mice were humanely euthanized according to guidelines established in our IACUC protocol by carbon dioxide inhalation and cervical dislocation.

#### Cdh5-Cre; TIE2^L914F^ mice

To generate experimental offspring, female *B6-Tg(Rosa26-TIE2^L914F^)^EBos^* mice were crossed with male *Tg(Cdh5-cre/ER^T2^)^1Rha^*(*iCdh5-Cre*) mice (36). For systemic induction of the transgene, offspring were administered 0.5µg tamoxifen (Sigma, cat# T5648, dissolved in sunflower oil) via intraperitoneal (i.p.) injection at postnatal day 5 (P5). Mice were then either monitored for survival or their tissues were collected at P23 for analysis of vascular parameters and immunostaining. For topical induction of the transgene in the time course model, offspring were administered 25µg of 4-hydroxytamoxifen (4-OHT) (Sigma, cat#H7904, dissolved in acetone) by dropping 5µl of stock 4-OHT (at a concentration of 5mg/ml) onto the ears of mice at P10 with a 20µl micropipette. Mice were then either monitored for survival or their ear tissues were collected at P17, P21, or P25 for immunostaining and analysis.

#### Drug administration

For topical induction of the transgene in the drug administration model, offspring were administered 5µg of 4-hydroxytamoxifen (4-OHT) (Sigma, cat# H7904, dissolved in acetone) by dropping 5µl of stock 4-OHT (at a concentration of 1mg/ml) onto the ears of mice at P10 with a 20µl micropipette. Drug dosing was initiated two days after tamoxifen induction (P12) and was continued every other day throughout the course of the experiment. Mice were either monitored for survival or their ear tissues were collected at the indicated collection timepoint. Longitudinal imaging of mouse ears was performed twice per week (every 3-4 days) beginning at P21.

For all experiments, *iCdh5-Cre* only littermate mice were used as controls for *Cdh5-CreER^T2^; TIE2^L914F^* mutant mice.

### Rapamycin experiments

Rapamycin (MedChem Express, cat# HY-10219) was dissolved in DMSO to make a 5mg/ml stock solution and then further diluted in vehicle to a concentration of 0.2mg/ml. Vehicle consisted of 5% PEG300 (Sigma, cat# 807484), 0.25% Tween80 (Fisher Scientific, cat# BP338-500), in saline (Baxter, cat#2F7124). Mice were then randomly sorted into rapamycin and vehicle groups that received 2mg/kg rapamycin i.p. (10µl per gram of mouse body weight) or vehicle with equivalent amounts of DMSO (4%).

#### iVEC-Cre; TIE2^L914F^ mice

To generate experimental offspring, female *B6-Tg(Rosa26-TIE2^L914F^)^EBos^* mice were crossed with male *Gm5127(BAC)CreER^T2^*(*iVEC-Cre*) mice(39). For topical induction of the transgene, offspring were administered 25µg of 4-OHT onto the ears of mice at P10. Mice were then monitored for formation of vascular lesions and their ear tissues were collected at P52 for immunostaining and analysis. *iVEC-Cre* only littermate mice were used as controls for *iVEC-Cre; TIE2^L914F^*mutant mice.

### Isolation of ear skin whole-mounts

Isolation of whole-mount skin tissues from dissected mouse ears was performed based on the protocol established in Yamazaki et. al. (2018) (53). Upon dissection, ears were fixed overnight at 4°C in 10% neutral-buffered formalin and then subsequently stored in PBS (Fisher Scientific, cat#BP3994). To prep the ear for whole-mount dissection, hair was removed by plucking with ultra-fine forceps (Fine Science Tools, cat# NC9100523). The inner ear skin was then carefully removed from the underlying cartilage, which was then subsequently stripped from the outer ear skin layer. The skin whole-mounts were then used for immunostaining, imaging, and analysis. For all analyses, the same skin layer (either outer or inner) was used for all mice included in that analysis.

### Histology and immunostaining

For histological imaging of tissues, tissues were fixed as above and then paraffin embedded and sectioned at 5µm for hematoxylin and eosin (H&E) staining.

For immunostaining of tissue sections, tissues were formalin fixed, paraffin embedded, and sectioned as above. Sections were then deparaffinized with xylene and rehydrated through a descending ethanol series. Antigen retrieval was performed by boiling the slides in 0.01 M citric acid (pH 6.0). Slides were blocked for 1 h at room temperature using the following antibody buffer: 5% bovine serum albumin (BSA, Sigma, cat#A7906) in PBS, containing 0.5% Triton X-100 (Sigma, cat#X100). Immunostaining was then performed with antibodies listed in **Table 2**. All primary antibodies were incubated overnight at 4°C in antibody buffer. If necessary, secondary antibodies (or streptavidin-fluorophore conjugates) were added the next day and incubated for 2 h at room temperature in antibody buffer. All sections were additionally counterstained with 4’,6-Diamidino-2-Phenylindole, Dihydrochloride (DAPI, cat#D1306, Invitrogen, 5 µg/mL) for 30 min at room temperature to visualize nuclei. Sections were mounted with ProLong™ Gold Antifade Mountant (Invitrogen, cat# P36930).

For immunostaining of whole-mount samples (embryos or ear skin), they were also blocked for 1 h at room temperature in antibody buffer (5% BSA 0.5% Triton X-100 in PBS) in the wells of a 24-well plate (Fisher, cat #08-772-1). Immunostaining was then performed as described above, except all incubations (including secondary antibodies and streptavidin-fluorophore conjugates) were performed overnight at 4°C. Whole-mounts were mounted with Fluoromount-G® mounting medium (Southern Biotech, cat#0100-01) and sealed with a coverslip.

Due to low signal of the mCherry reporter from tissue processing and paraffin embedding, protein expression was detected by immunostaining and tyramide amplification. For detection in tissue sections, samples were first incubated in 0.3% hydrogen peroxide for 15 minutes to quench endogenous peroxidases. For detection in ear skin whole-mounts, samples were incubated in 3% hydrogen peroxide for 1h. All samples were then blocked/permeabilized in antibody buffer for 1 h. Anti-RFP (mCherry) primary antibody was then incubated overnight at 4°C in antibody buffer, followed by anti-Rabbit HRP for 2h at room temperature (tissue sections) or overnight at 4°C (whole-mounts). Fluorescent detection was performed using CF® 594-tyramide conjugates (Biotium, cat# 92174) according to the manufacturer’s instructions. Immunostaining for additional markers of the vasculature was then performed as described previously.

### EdU detection of cell proliferation in vivo

To label proliferating cells in mice, we utilized incorporation of 5-ethynyl-2’-deoxyuridine (EdU) into cells undergoing DNA replication. EdU was dissolved in PBS to prepare a stock solution at 5mg/ml and then injected intraperitoneally at 50mg/kg, 24h before tissue harvest. Tissue samples (sections or whole-mounts) were then prepared as described previously. Prior to any other immunostaining, EdU incorporation was detected using the Click-iT™ EdU Alexa Fluor™ 647 dye kit (Invitrogen, cat# C10340), according to the manufacturer’s instructions. Samples were then further immunostained for endothelial cell nuclei and other vascular markers.

### Confocal imaging and quantification

Z-stack images of tissue sections and whole-mounts were acquired on a Nikon A1 Inverted LUNV laser-scanning confocal microscope in the CCHMC Bio-Imaging and Analysis Facility (RRiD SCR_022628). Nikon NIS Elements software was used to create maximum intensity projections of the images.

#### Tissue Section Vascular Parameters Quantification

Quantification of various parameters of the vasculature (vessel density, vascular area, vessel size distribution) in tongue and tail sections was performed using Fiji (ImageJ)(54) software. Vessels were identified by thresholding of CD31 signal, which were manually verified to ensure that vessel lumens were included in the measured areas. Vessel density was calculated as the number of vessels in the total tissue area. Vascular area was calculated as the total area covered by vessels/vessel lumens over the total tissue area. Individual vessels were also stratified by size: small less than 500µm^2^, medium between 500µm^2^ and 1000µm^2^, and large greater than 1000µm^2^. A minimum of 150 vessels were analyzed per mouse to yield a representative average value.

#### αSMA Coverage Quantification

Ear skin whole-mounts from mutant mice were immunostained for CD31 and αSMA. Non-lesional and lesional veins were identified by morphology and vessel size. Using Fiji (ImageJ) software, CD31 staining of vessels was used to create a region of interest (ROI) encompassing the vessel area. Separately, a threshold of αSMA staining was used to identify positive areas. The vessel ROI was then used as a mask over the αSMA threshold to measure the percent positive αSMA area of the vessel. Results are shown as αSMA+ vessel area over total vessel area. A minimum of 8 vessels of each type were analyzed and averaged per mouse.

#### EdU Quantification

Quantification of EdU+ EC in ear skin whole-mounts was performed using the General Analysis 3 tool in NIS Elements software, as we have described previously (55). In short, images were first masked on staining of the relevant vasculature (Endomucin or CD31). Within this mask, the number of relevant EC nuclei (labeled by ERG, SOX17, or PROX1 immunostaining) were counted using the “BrightSpots” function, to yield the total number of cells. Then, the EC nuclear signal was used as a mask for the EdU signal, isolating nuclei that were positive for both. These signals were then also counted using the “BrightSpots” function, which yielded the number of double-positive (EdU+ nuclear marker+) nuclei. The number of proliferating cells within the relevant vasculature was then represented as the number of double-positive nuclei over the total number of nuclei. 5 images were analyzed and averaged per mouse.

Quantification of EdU+ EC in tissue sections was performed using Fiji (ImageJ) software. For control animals, the number of EdU+ ERG+ Endomucin+ nuclei were manually counted and then represented as a total of ERG+ Endomucin+ nuclei. For mutant animals, lesional vessels were defined as having diameters larger than 30µm. Specifically within lesional vessels, the number of EdU+ ERG+ Endomucin+ nuclei were counted in the same manner as in the control animals. 4-11 images were analyzed and averaged per mouse.

#### Cell Size Quantification

Quantification of cell size was performed using Fiji (ImageJ) software. Within ear skin whole-mounts of mutant animals, Endomucin+ non-lesional and lesional veins were identified by size and morphology. Using CD31 as marker of EC borders, individual cells were manually outlined in Fiji and their area was used as a proxy of cell size. For each animal, 50-300 cells were measured across 3-8 individual vessels.

### Longitudinal vascular lesion quantification

In the rapamycin treatment studies, lesion formation in the ears was monitored on live mice by longitudinal imaging twice per week, beginning at P21. Mice were anesthetized using isoflurane and their ears were imaged with a Leica S9 D Stereomicroscope (cat# 10450815) equipped with a 0.63x lens (cat# 10450818) and a Flexacam C3 camera (cat # 12730533).

Quantification was then performed using Fiji (ImageJ) software, in which lesions were manually outlined. Lesion area was then normalized to total ear area. For each mouse, the values for both ears were averaged to yield a single value at each timepoint.

### Mouse EC isolation and immunoblotting

3-week old mice were euthanized by CO_2_ inhalation and cervical dislocation. Their tails were removed and the soft tissue was cut longitudinally down the length of the tail, allowing the tail bones to be carefully removed. Then, the soft tissue was incubated in a 1U/ml solution of dispase in PBS (StemCell, cat #07913) for 45 minutes at 37°C on a rocker. For each tail, 10ml of digestion solution was prepared: Dulbecco’s modified Eagle medium (DMEM) (Gibco, cat# 11965118) with 2% fetal bovine serum (FBS) (Hyclone, cat# SH30910.03), 1.26mM Ca^2+^, 0.81 mM Mg^2+^, 1 mg/ml of collagenase A (Roche, cat# 10103578001), and 7.5µg/ml DNAse (Sigma, cat# D4527).

After digestion in dispase, the epidermis was carefully removed from the underlying dermal tissue, which was then finely minced using scissors and placed in 10ml of digestion solution for 45 minutes at 37°C on a rocker. After digestion, the tissue was triturated with a micropipette until no chunks were remaining. The solution was filtered through a 70µm filter, which was then washed 3x with PEB buffer (2mM EDTA, 0.5% BSA, in PBS). The cell suspension was then centrifuged at 2100 rpm for 10 minutes to pellet the cells. The cell pellet was resuspended in 1ml of RBS lysing buffer (Sigma, cat# R7757) for 3 minutes. 10ml of PEB was added to dilute the RBC lysing buffer and then centrifuged at 1500 rpm for 5 min. Samples were then processed using MACS CD31 microbeads (Miltenyi, cat# 130-097-418) and the OctoMACS separator (cat# 130-042-109) with MS columns (cat# 130-042-201) to isolate CD31+ ECs, according to the manufacturer’s protocol.

Isolated mouse EC were cultured in vitro on gelatin-coated (BD Difco, cat# 214340) plates and cultured in endothelial growth medium (EGM2, Lonza, cat# CC-3162) supplemented with 10% FBS (Hyclone, cat#SH30910.03) and 1% penicillin-streptomycin-glutamine (Corning, cat# 30-009-CI). Upon reaching 80% confluency, cells were treated with 2µM 4-OHT in 2% FBS EGM2 media to induce transgene expression and then collected for protein 48h later. For rapamycin treatment, cells were treated with 15nM rapamycin in DMSO concurrently with 4-OHT induction and were collected for protein 48h later.

For protein collection, cells were washed with PBS then lysed using ice-cold RIPA lysis buffer (Boston Bioproduct, cat# BP-115-5) supplemented with HALT^TM^ protease/phosphatase inhibitor cocktail (Thermo Scientific, cat# 78442). The protein concentration was determined using the BCA Protein Assay Kit (Thermo Scientific, cat# PI23225). 30 µg of total protein were subjected to SDS-PAGE (4-20%, Midi Criterion precast gels, Bio-Rad, cat#5678094) and transferred to a PVDF membrane (Immobilon®-P PVDF Membrane, Millipore, cat#IPVH00010). Membranes were blocked in 5% nonfat dried milk for 1 h at room temperature and probed with the antibodies listed in **Table 2** in buffer consisting of 2% BSA in TBST, overnight at 4°C. Membranes were then incubated with secondary antibodies in 5% nonfat dried milk, for 2 h at room temperature. For HRP secondaries, antigen–antibody complexes were visualized using Classico HRP chemiluminescent substrate (Sigma, cat# WBLUC0500) on a ChemiDoc imager (BioRad). For fluorescent secondaries, antigen–antibody complexes were visualized using Odyssey Scanner and analyzed using Image Studio Software (LI-COR, v5.2). Densitometric quantification of western blot bands was performed using Image Lab (BioRad) software. GAPDH was used as a loading control for all blots.

### Data collection and statistics

Prism 9.0 software (GraphPad) was used to make graphs and perform statistical analysis. Data are presented as mean ± standard deviation or mean ± standard error of the mean (as indicated in figure legends) with n values reported in figure legends. Statistical significance between 2 groups was assessed by unpaired Welch’s *t*-test or by ordinary one-way ANOVA with Tukey’s multiple comparisons for analyses with 3 or more groups. Statistical significance between groups with repeated measurements (such as weight or lesion formation over time) were analyzed with two-way ANOVA. Statistical significance is defined as *p* values less than 0.05. Schematics were created with Biorender.com.

## Study Approval

All mouse experiments and procedures have been reviewed and approved by the CCHMC Institutional Animal Care and Use Committee (protocol number IACUC 2023-0025 and 2026-0023). All animals were cared for in accordance with guidelines from the National Institutes of Health. Studies were performed using both male and female mice and data analyzed as combined and separately.

## Data Availability

The data that support the findings of this study are presented here and/or available from the corresponding author upon reasonable request.

## Acknowledgements and sources of funding

Research reported in this manuscript was supported by the National Heart, Lung, and Blood Institute, under Award Numbers R01HL117952 (E.B.), R01HL167700 (E.B.), and F31HL176101 (L.J.B.), part of the National Institutes of Health. The content is solely the responsibility of the authors and does not necessarily represent the official views of the National Institutes of Health. This manuscript is the result of funding in whole or in part by the National Institutes of Health (NIH). It is subject to the NIH Public Access Policy. Through acceptance of this federal funding, NIH has been given a right to make this manuscript publicly available in PubMed Central upon the Official Date of Publication, as defined by NIH. Additional funding supporting the study was provided by the American Heart Association (AHA) Predoctoral Fellowship (https://doi.org/10.58275/AHA.24PRE1191403.pc.gr.190588) to L.J.B. We thank the Bio-imaging and Analysis Facility and the Comparative Medicine Division at Cincinnati Children’s Hospital Medical Center for providing state-of-the-art instrumentation, services, training, and education.

## Author contributions

L.J.B, S.S., and E.B. conceived the project. E.B. supervised the research. M.S. provided Gm5127CreER^T2^ mice and guidance on data interpretation and analysis. L.J.B., S.S., K.S., and C.S. performed the mouse experiments and sample collection. L.J.B. and A.B. performed immunostaining and confocal imaging. L.J.B., S.S. and A.B. performed quantifications. L.J.B. wrote the manuscript draft and prepared the figures. All authors revised and commented on the manuscript.

