## Supplementary Data for "An endothelial-specific TIE2-L914F mouse model reveals cellular mechanisms of venous malformation progression and recapitulates the patients’ response to targeted therapy"

**Content:**

Supplementary Figures S1-S9

Supplementary Tables S1 and S2.

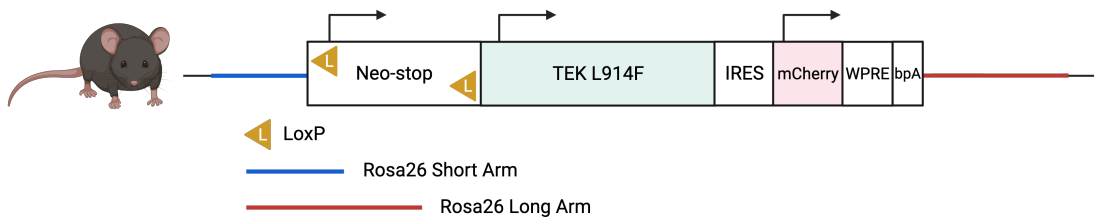

**Supplementary Figure S1: Transgenic construct of *B6-Tg(Rosa26-TIE2<sup>L914F</sup>)EBoS* mice.** TIE2<sup>L914F</sup> were generated by insertion of an inducible *TEK* mutant construct at the Rosa26 genomic locus. The *TEK* human transgene, containing the p.L914F mutation, is controlled by a floxed Neo-stop cassette. It is followed by an IRES (internal ribosome entry site) sequence that allows for co-expression of an mCherry fluorescent reporter. WPRE = Woodchuck Hepatitis Virus posttranscriptional regulatory element. bpA = bovine polyadenylation signal.

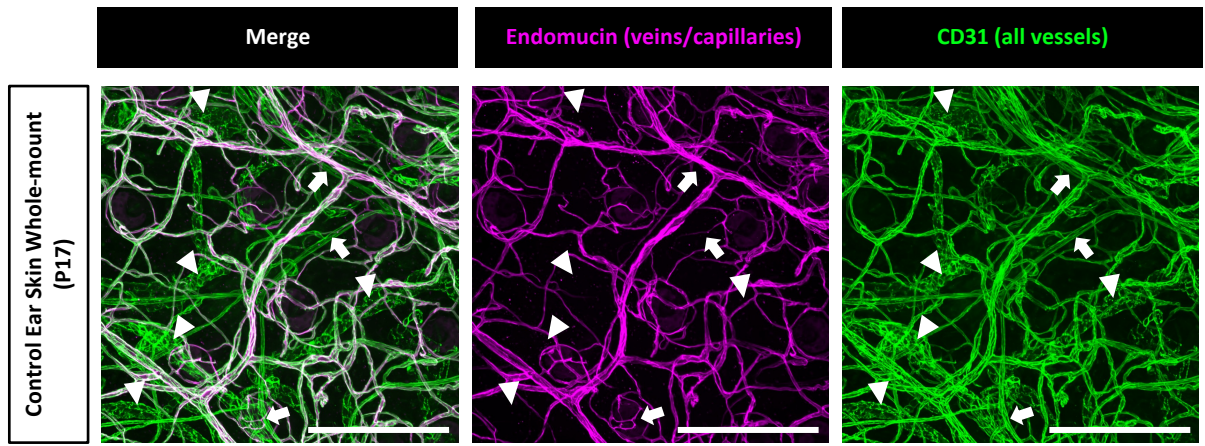

**Supplementary Figure S2: Endomucin immunostaining is specific for veins/capillaries in P17 ear skin whole-mounts.** Whole-mount ear skin from wild-type mouse was immunostained for Endomucin (magenta) and CD31 (green). Arrows indicate Endomucin-negative arterial vessels. Arrowheads indicate Endomucin-negative lymphatic vessels, which can be distinguished by their dilated morphology and discontinuous, button-like expression of CD31. In the merged image, veins/capillaries appear white due to the overlap of Endomucin and CD31 signal. Scale bar = 250μm.

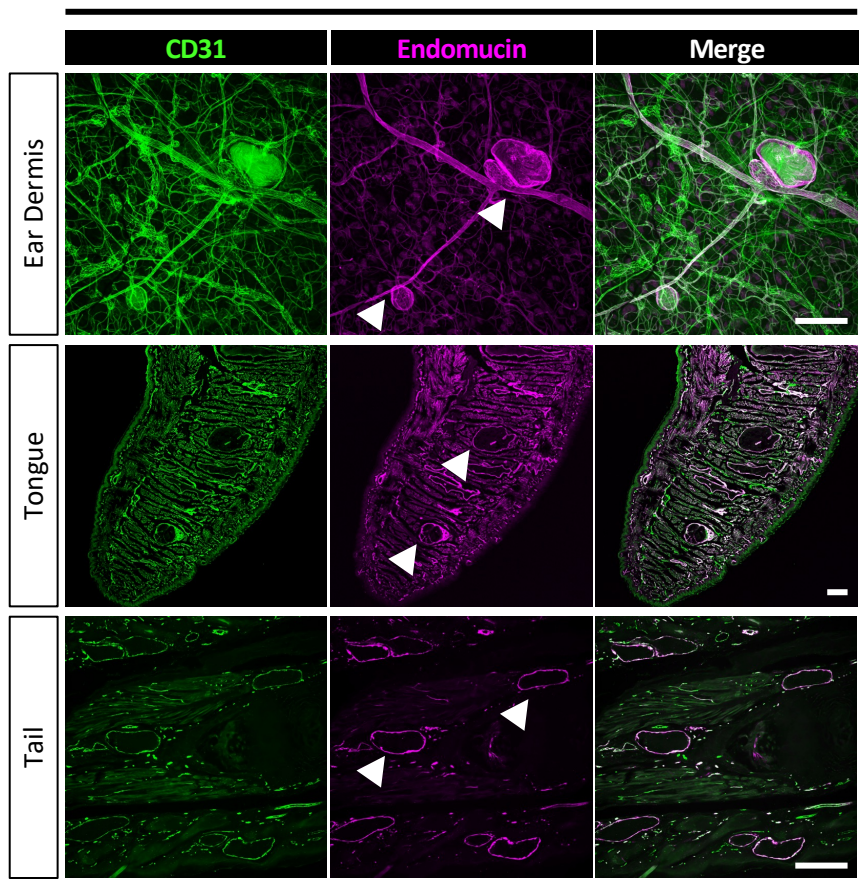

**Supplementary Figure S3: Abnormally dilated blood vessels in iCdh5-TIE2<sup>L914F</sup> mice treated with systemic tamoxifen are positive for Endomucin, a marker of venous/capillary vessels.** Mice and tissue samples are from the experiments performed in Figure 1. Whole-mounts (top row, ear skin) or tissue sections (bottom rows, tongue and tail) were immunostained for CD31 (green, pan-vasculature) and Endomucin (magenta, veins/capillaries). Scale bar = 250μm.

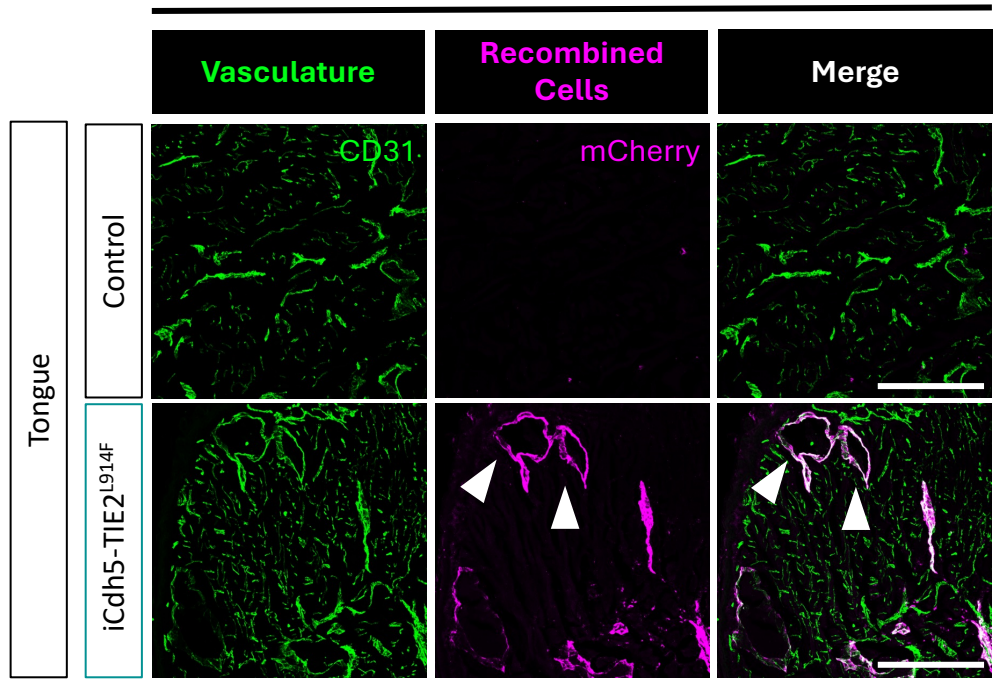

**Supplementary Figure S4: Abnormally dilated blood vessels in iCdh5-TIE2<sup>L914F</sup> mice treated with systemic tamoxifen are lined by mCherry+ mutant EC.** Mice and tongue tissue samples are from the experiments performed in Figure 1. Tongue sections were immunostained for CD31 (green, pan-vasculature) and mCherry (magenta, recombined cells). Scale bar = 250μm.

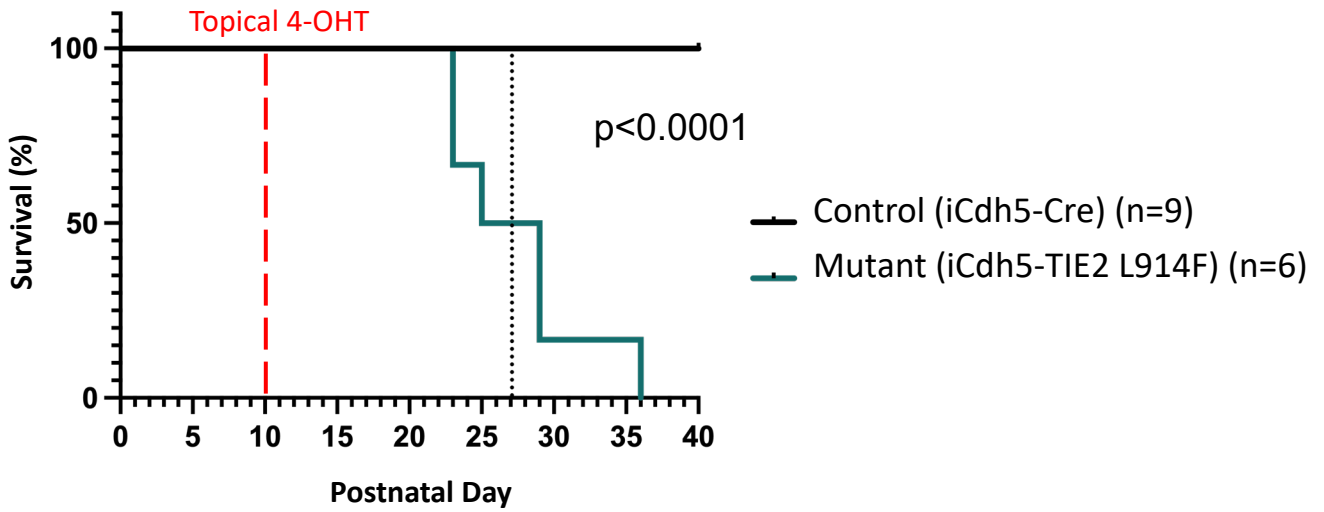

**Supplementary Figure S5: Survival curve of mice from the topical time course model.** Kaplan-Meier survival curve of control (black line) and mutant mice (teal line) treated with 25μg of 4-OHT on P10. The vertical dotted line indicates 50% survival of mutant mice. Gehan-Breslow-Wilcoxon test ($p < 0.0001$).

### Whole Mount Ear Skin from Wild-Type P17 Mice

Masked on EMCN or CD31

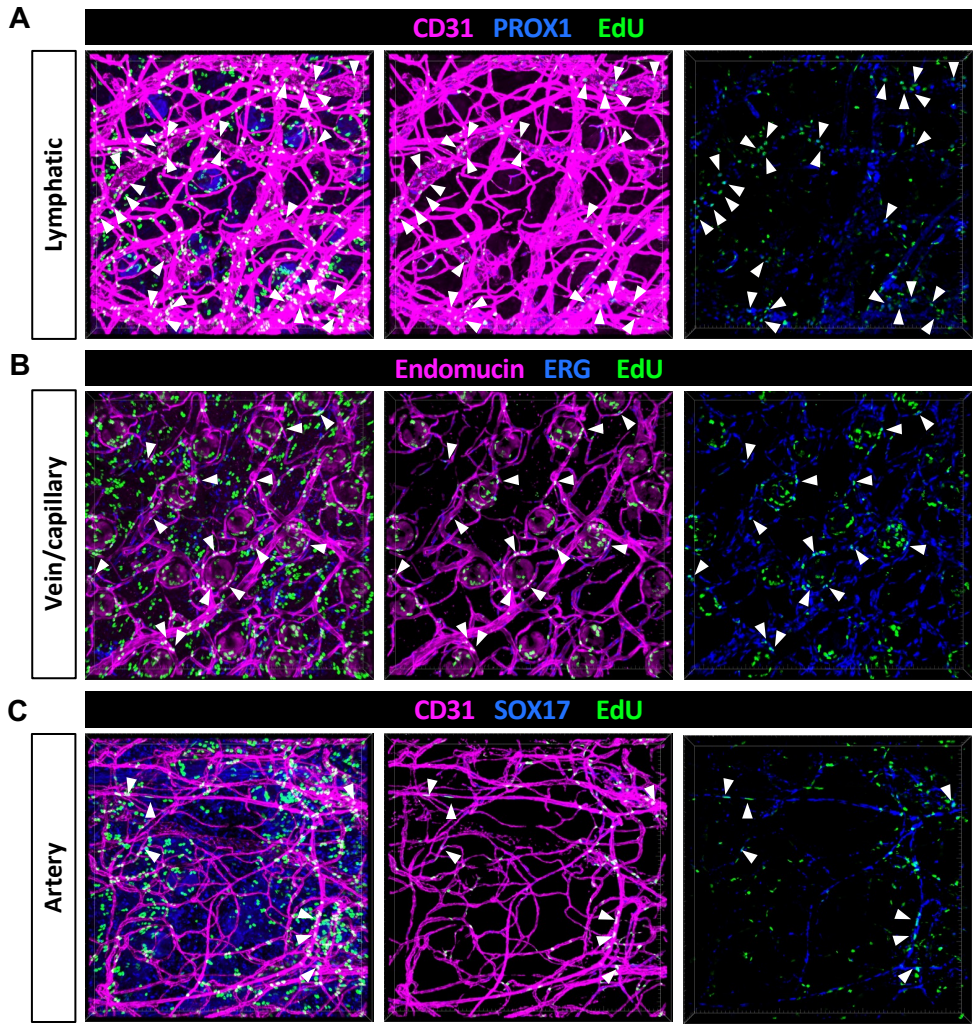

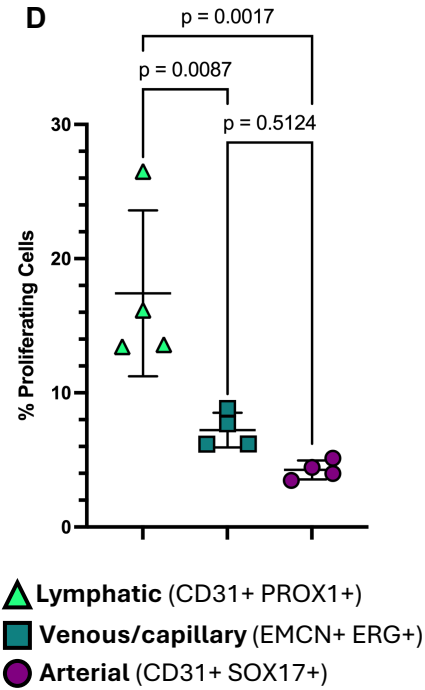

**Supplementary Figure S6: EC proliferation in various vessel types at baseline.** (A) Whole-mount ear skin tissues were isolated from wild-type (WT) mice at P17, that were previously treated with EdU 24h prior to label proliferating cells. Lymphatic EC proliferation was found by staining for EdU positive (green) lymphatic EC nuclei (PROX1, blue) within CD31 positive vessels (magenta). (B) Endomucin (magenta) and ERG (blue) were used to identify EC nuclei within the venous/capillary plexus. (C) Arterial nuclei (SOX17, blue) were identified in CD31 positive vessels (magenta). Masking on EMCN or CD31 alone includes some non-EC proliferating cells due to autofluorescence of hair follicles (right-most panels). Using a secondary EC nuclear stain (PROX1, ERG, or SOX17) further excludes any non-EC signal. (D) Using Nikon General Analysis 3 software, the number of total nuclei within each vessel type was calculated (lymphatic – CD31+/PROX1+; venous/capillary – Endomucin+/ERG+; arterial – CD31+/SOX17+). The number of these nuclei that were also positive for EdU was then calculated and the number of proliferating cells is represented as the percentage of EdU nuclei over total nuclei. Mean±SD. 1way ANOVA with Tukey's multiple comparisons.

**A**

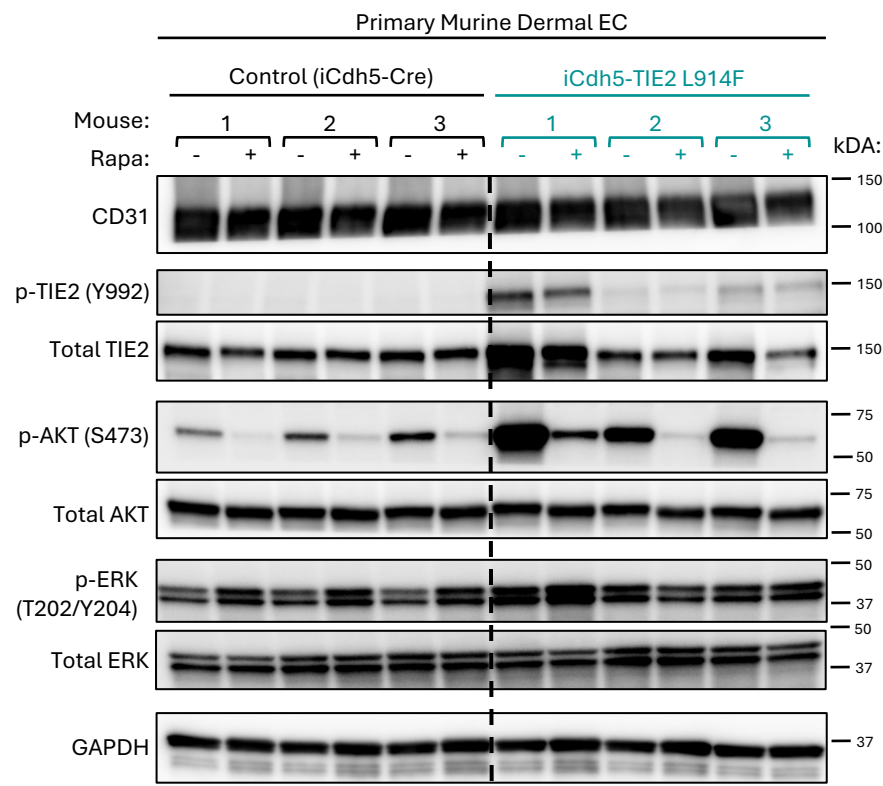

**B**

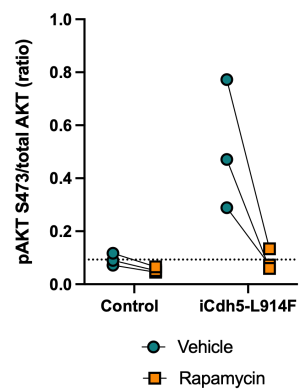

**C**

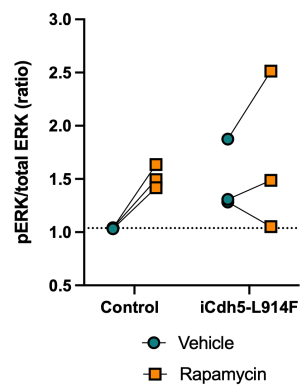

**Supplementary Figure S7: Rapamycin reduces AKT activation in dermal EC isolated from iCdh5-TIE2<sup>L914F</sup> mice.** (A) Primary EC were isolated from tail dermis via magnetically activated cell sorting, cultured and induced with 4-OHT in vitro, and analyzed with western blot for the indicated proteins. Cells were collected from 3 individual control and 3 mutant mice. Cells from each mouse were split and treated with vehicle or 15nM rapamycin for 48h before collection. (B) Densitometric analysis of pAKT and (C) pERK. Results are shown as the ratio of phospho-protein to total protein. Paired lines connect cells from the same mouse treated with vehicle or rapamycin. Dotted lines indicate the average value of the vehicle-treated cells from control mice.

**A**

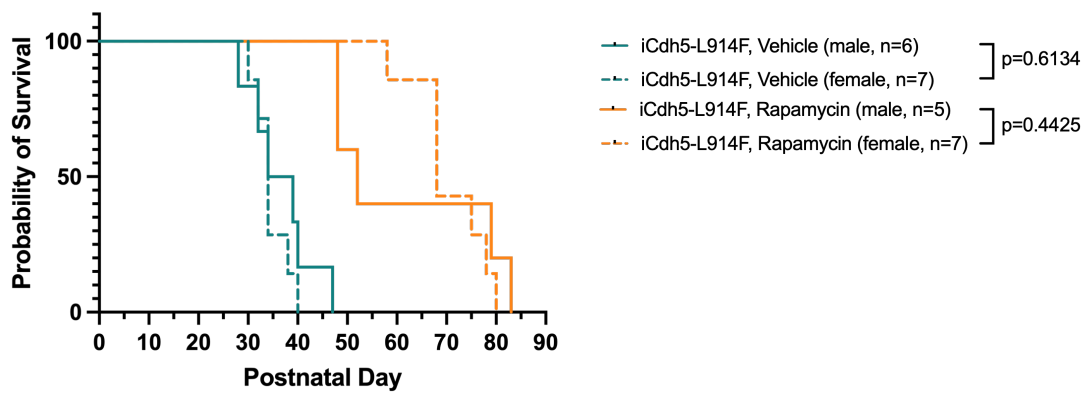

**B**

Male

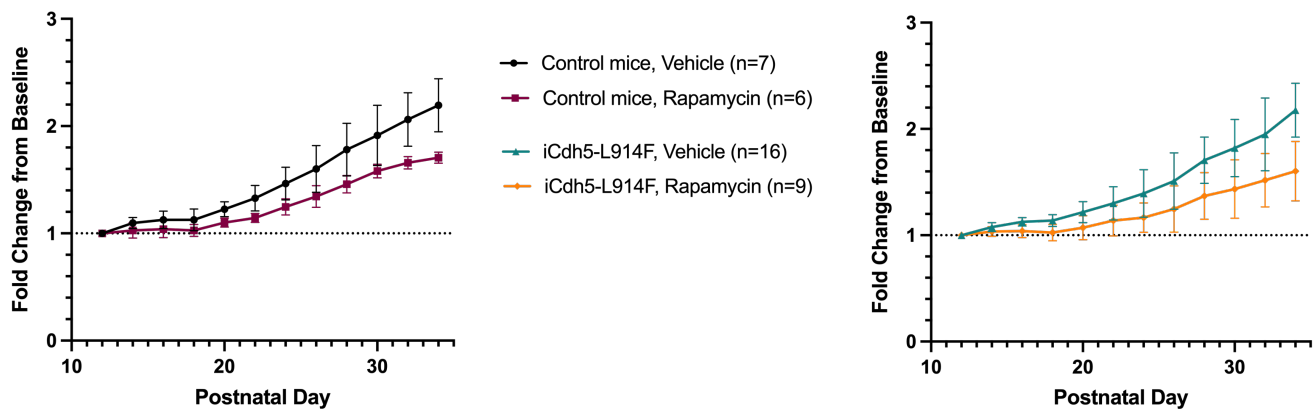

**C**

Female

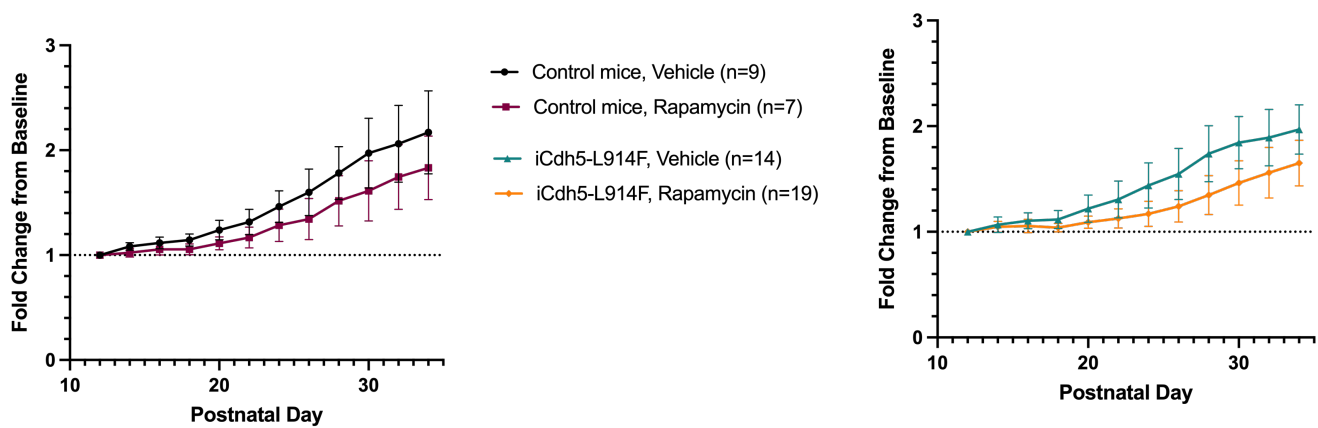

**Supplementary Figure S8: Rapamycin-treated mouse sex-based survival and weight curves.** (A) Survival curve from Figure 6D with each mutant group separated into male and female categories. p-values of inter-group comparisons (male vs female) is shown in the figure legend (Gehan-Breslow-Wilcoxon test). (B) Weight data from all male mice (collection and survival cohorts). Control (left) and mutant (right) mice treated with vehicle or rapamycin were weighed every other day beginning at P12. (C) Weight data from female mice.

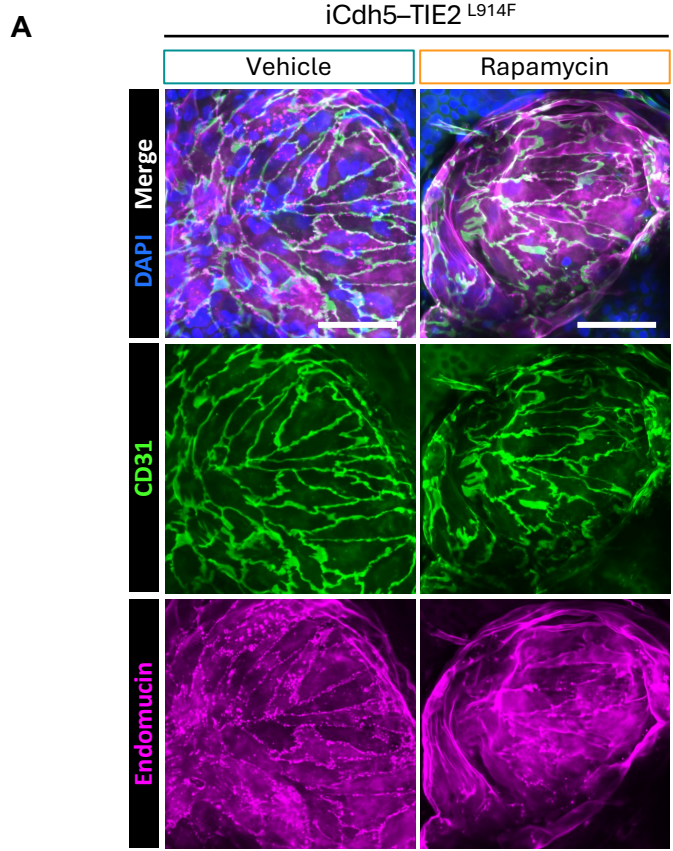

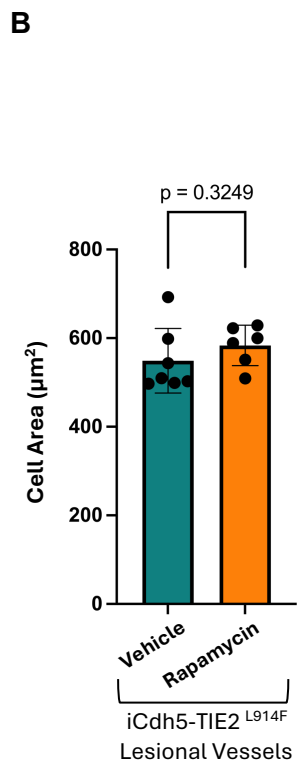

**Supplementary Figure S9: Rapamycin treatment does not rescue EC enlargement in iCdh5-TIE2<sup>L914F</sup> mice.** (A) iCdh5-TIE2<sup>L914F</sup> mice treated with rapamycin or vehicle (from Figure 6) were analyzed for lesional EC size. Ear skin whole-mounts were immunostained for Endomucin (magenta, veins/capillaries) and CD31 (green, EC cell borders). Scale bar = 50µm. (B) CD31+ cell borders were manually outlined to quantify cell area of EC within lesional vessels. n=6-7 mice per group. Mean±SD. Welch's t-test.

#### Supplementary Table 1: Genotyping primers.

| Mouse Line: | Forward (5'-3') | Reverse (5'-3') | 2<sup>nd</sup> Reverse (if applicable) |
| --- | --- | --- | --- |
| **TIE2<sup>L914F</sup>** | AGC ACT TGC TCT CCC AAA GTC | AAT CTG TGG GAA GTC TTG TCC | CCA GAG GCC ACT TGT GTA GC |
| **Cdh-Cre<sup>ERT2</sup>** | GCG GTC TGG CAG TAA AAA CTA TC | GTG AAA CAG CAT TGC TGT CAC TT |  |
| **CMV-Cre** | ACCAGCCAGCTATCAACTCG | TTACATTGGTCCAGCCACC |  |
| **mTmG** | CTC TGC TGC CTC CTG GCT TCT | CGA GGC GGA TCA CAA GCA ATA | TCA ATG GGC GGG GGT CGT T |
| **iVEC-Cre** | GCA TCT TTA TCC TCC CTG GAC | CCT GTT TTG CAC GTT CAC CG |  |

#### Supplementary Table 2: List of antibodies.

IF = immunofluorescence; WB = Western blot, CST = Cell Signaling Technology

| Host Species | Antigen | Conjugate | Company | Catalog Number | Clone | Dilution | Stock Concentration | Application |
| --- | --- | --- | --- | --- | --- | --- | --- | --- |
| Primary Antibodies |  |  |  |  |  |  |  |  |
| Goat | CD31 |  | R&D | AF3628 | N/A | 1:100 | 0.2 mg/ml | IF |
| Goat | CD31 | Alexa 488 | R&D | FAB3628G | N/A | 1:100 | 0.05 mg/ml | IF |
| Rabbit | mCherry |  | Rockland | 600-401-379 | N/A | 1:100 | 1 mg/ml | IF |
| Rat | Endomucin |  | Santa Cruz | sc-65495 | V.7C7 | 1:50 | 0.2 mg/ml | IF |
| Rabbit | ERG |  | Abcam | AB92513 | EPR3864 | 1:100 | 0.9 mg/ml | IF |
| Goat | SOX17 |  | R&D | AF1924 | N/A | 1:50 | 0.2 mg/ml | IF |
| Goat | PROX1 |  | R&D | AF2727 | N/A | 1:50 | 0.2 mg/ml | IF |
| Mouse | αSMA | FITC | Sigma | F3777 | 1A4 | 1:100 | 2.7 mg/ml | IF |
| Rat | LYVE1 |  | R&D | MAB2125 | 223322 | 1:50 | 0.5 mg/ml | IF |
| Mouse | AKT |  | CST | 2920 | 40D4 | 1:500 | 27 µg/ml | WB |
| Rabbit | p-AKT T308 |  | CST | 13038 | D25E6 | 1:500 | 613 µg/ml | WB |
| Rabbit | p-AKT S473 |  | CST | 4060 | D9E | 1:1000 | 91 µg/ml | WB |
| Rabbit | p-ERK 1/2 T202/Y204 |  | CST | 4370 | D13.14.4E | 1:500 | 502 µg/ml | WB |
| Mouse | ERK1/2 |  | CST | 4696 | N/A | 1:500 | 251 µg/ml | WB |
| Mouse | TIE2 |  | CST | 4224 | AB33 | 1:500 | 543.8 µg/ml | WB |
| Rabbit | p-TIE2 Y992 |  | R&D | AF2720 | N/A | 1:200 | 0.2 mg/ml | WB |
| Mouse | GAPDH |  | Millipore | MAB374 | 6C5 | 1:5000 | 1 mg/ml | WB |
| Secondary Antibodies |  |  |  |  |  |  |  |  |
| Goat | α-Rabbit IgG | Alexa 488 | Invitrogen | A32731 | N/A | 1:200 | 2 mg/ml | IF |
| Goat | α-Rabbit IgG | Alexa 647 | Invitrogen | A21244 | N/A | 1:200 | 2 mg/ml | IF |
| Donkey | α-Goat IgG | Alexa 594 | Invitrogen | A32758 | N/A | 1:200 | 2 mg/ml | IF |
| Goat | α-Rat IgG | Biotin | Vector | BA-9401 | N/A | 1:200 | 0.5 mg/ml | IF |
|  | Streptavidin | DyLight 594 | Vector | SA-5594 | N/A | 1:200 | 1 mg/ml | IF |
|  | Streptavidin | DyLight 649 | Vector | SA-5649 | N/A | 1:200 | 1 mg/ml | IF |
| Goat | α-Rabbit IgG | HRP | Vector | PI-1000-1 | N/A | 1:200/<br>1:5000 | 1 mg/ml | IF/WB |
| Horse | α-Mouse IgG | HRP | Vector | PI-2000 | N/A | 1:5000 | 1 mg/ml | WB |
| Donkey | α-Mouse IgG | DyLight 680 | Invitrogen | SA5-10170 | N/A | 1:2500 | 0.5mg/ml | WB |
| Goat | α-Rabbit IgG | DyLight 800 | Invitrogen | SA5-10036 | N/A | 1:2500 | 0.5mg/ml | WB |
